# Generating *E. coli* 0.5 controlled by a half-sized genome

**DOI:** 10.64898/2026.06.01.729178

**Authors:** Takahito Mukai, Aika Ohishi, Emi Hagiuda, Keita Shimamoto, Koki Yoshida, Masayuki Su’etsugu

## Abstract

Genome synthesis is a major limitation in generative biology. Here, the half-sized genome of *Escherichia coli* was constructed by fleshing out an imperfect minimal genome through genome-scale debugging process. Our platform consists of integrated development environment (IDE) and runtime environment (RTE). The genome IDE supported the cell-free assembly of 200–300 kb plasmids and their in vivo fusion into a single 1.7 Mb plasmid. This imperfect genome was stably maintained in *E. coli* as a guest genome. The RTE relies on the restriction enzyme-mediated self-digestion of the host genome in the presence and absence of the RecA recombinase. The guest genome was tested, debugged, and partially replaced by the host genome to establish *E. coli* controlled by a 2.3-Mb genome. This is less than half in size of the wildtype and the smallest ever reported. Enfleshing a guest genome will facilitate genome printing that transforms AI-designed genomes into physical ones.

## Introduction

In the era of generative biology, synthetic genomes will be designed by genome-generative AI (*1–5*) and transformed into physical genomes via genome printing. However, there is a chicken and egg situation. It is challenging for AI to design unprecedented things such as the minimal genome of *Escherichia coli*. The genome printing technology is also underdeveloped for the rapid prototyping of bacterial genomes. It is required to develop an actual platform to generate and implement a genome instance. Otherwise, there will be no data for AI to learn.

Since a half century ago (*6*), molecular cloning technology has widened the range of applications. In the last two decades, *Saccharomyces cerevisiae* has been routinely used for the assembly and cloning of eukaryotic chromosomes and small bacterial genomes (*7–10*). The Sc2.0 consortium established a yeast strain with a half-synthetic genome through the debug and consolidation of the synthetic chromosomes (*11–14*). Bacterial mycoplasma genomes have been engineered because they are small enough to transplant into cells (*8–10, 15, 16*). In sharp contrast, molecular cloning of common bacterial genomes has been hampered by their large genome size (*17–19*). The whole genome of *E. coli* was replaced by the iterative replacement of the original genome segments with synthetic ones (*20–22*). However, this template-dependency has limited the design flexibility (*18, 23*). One possible solution would be the stepwise assembly of a guest genome for the subsequent genome swapping (*5*).

In the last few years, *E. coli* becomes a useful tool for megabase-sized chromosome engineering and a good alternative to yeast in the genome synthetic biology field (*19, 23–28*). Modernization of the traditional molecular cloning tools and the use of CRISPR/Cas enabled the fission and fusion of the genome and the assembly and cloning of a one megabase (1 Mb) chromosome (*25–31*). Genome swap is facilitated by establishing multipartite-genome strains of *E. coli* (*28, 30, 31*). Enzymatic supercoiling and repair (SCR) converts extracted chromosomes into compact molecules resistant to in vitro manipulation (*31*). Anucleate *E. coli* cells or minicells may be a good chassis of a designer genome (*32, 33*). Almost escape-free production of anucleate cells (SimCells) was achieved by the tightly controlled expression of I-CeuI mega-restriction enzyme (*34, 35*). Although our understanding of the *E. coli* genome is far from complete, the EcoCyc database (*36*) serves as a good comprehensive reference. Altogether, it is straightforward to assemble, clone, run, and debug a designer genome of *E. coli* using *E. coli*.

## Results

### Kernel genome project

In this study, we aim at the molecular cloning of a size-reduced genome of *E. coli* using *E. coli* (Fig. 1). We designed a ‘kernel’ genome named after the kernel (the core programs) of a computer operating system. The prototype kernel genome was designed to be hopefully as follows: 1) small (1.7 Mb) but not minimal (*17, 37*); 2) cloneable as a guest genome and standalone as the sole genome; 3) capable of replication in vitro by using the replication cycle reaction (RCR) (*38*); 4) no site for AvrII restriction enzyme (C|<u>CTAG</u>G). We designed an integrated development environment (IDE) for the kernel genome (Fig. 1). The genome IDE includes both in vitro assembly and in vivo assembly systems. Our IDE should be compatible with CRISPR/Cas-based systems because none of them is used. Cloning of bacterial artificial chromosomes (BACs) using RecA-deficient *E. coli* facilitates the next-generation sequencing (NGS) analysis and the phenotype analysis of the assembled DNA chunks. We then designed a cellular runtime environment (RTE) for the kernel genome (Fig. 1). The host *E. coli* genome can be eliminated by expressing AvrII, and the guest genome will be subjected to a run test to revive the cell. In debug mode, RecA homologous recombination might generate chimeras between the host and guest genomes. Chimeras can be developed by conjugative recombination, too. The project’s data are listed in Supplementary tables S1 to S5 and provided as Supplementary files.

**Figure 1.**
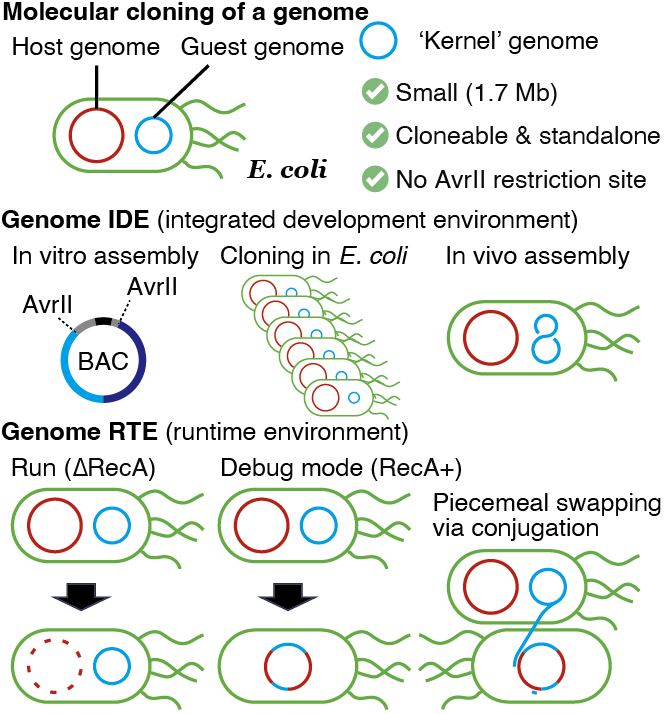
Plan of our *E. coli* kernel genome project. We aim at the molecular cloning of a genome in *E. coli* by developing a genome integrated development environment (IDE) and a genome runtime environment (RTE).

### Designing a kernel genome

We designed the prototype kernel genome based on the reduced genome of *E. coli* DGF-298W (*39*). While the wildtype *E. coli* K-12 strain W3110 has a 4.6 Mb genome, DGF-298W and its derivative RGF (*30*) have a 3 Mb genome. The prototype kernel genome (1.7 Mb) (Fig. 2A) inherits about half of the RGF genome to synthesize all the essential compounds and gains several wildtype loci that should be added back. In almost all cases, genes and operons are arranged without changing their relative locations of the wildtype genome, which may facilitate the RecA-mediated debugging. The genomic AvrII cut sites were totally eliminated. Since most of the 16S ribosomal RNA genes and all tRNA^Glu^ genes have a single AvrII cut site within their secondary RNA structures, the CCUAGG sequence was mutated by RNA engineering (Fig. 2A). The 16S RNAs were engineered to have the same local structure as the ΔCCUAGG 16S rRNAs, whereas the T-stem of tRNA^Glu^ was mutated without changing the affinity to EF-Tu according to the literature (*40*). Elimination of the genomic 16S rRNA and tRNA^Glu^ genes in *E. coli* SQ171 strain (*41*) was fully complemented with a plasmid expressing the ΔCCUAGG variants of 16S rRNA and tRNA^Glu^ from a modified *rrnC* operon. No read sequence for the wildtype genes was detected by the NGS analysis of the developed *E. coli* strain SQ171/pMW118cat-rrnCΔAvrII/ptRNA100-gltT::zeo.

**Figure 2.**
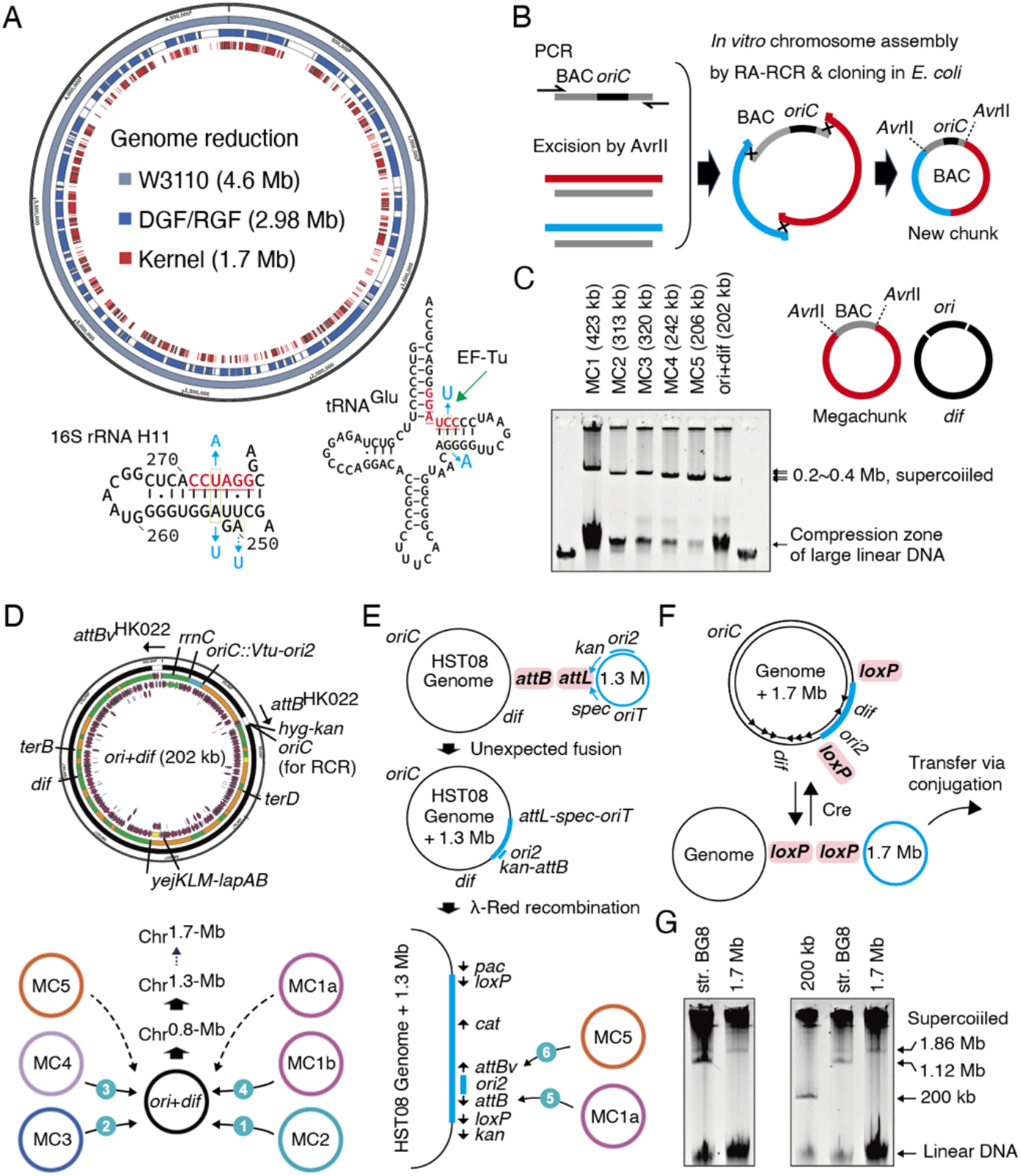
The integrated development environment for building the prototype kernel genome. **(A)** Designing the prototype kernel genome. Genome size reduction from the parental *E. coli* strains is depicted. Safe elimination of the AvrII sites in 16S rRNA genes and the tRNA^Glu^ genes is also depicted. **(B)** Scheme for the in vitro chromosome assembly followed by *E. coli* cloning. A BAC vector backbone containing the *oriC* sequence was prepared via PCR using primers containing a new AvrII site and the overlapping tail for a chunk to be assembled. Chunk DNAs were prepared by the AvrII digestion of chunk-containing plasmids lacking *oriC*. **(C)** The *E. coli* cloning of the five megachunks (MCs) and the *ori* and *dif* regions. The five megachunk plasmids and the *ori-dif* fusion plasmid were purified by miniprep and resolved by canonical agarose gel electrophoresis. **(D)** The *ori-dif* chromosome serves as the core to be equipped with the megachunks in the indicated order using the site-specific integrase of HK022 phage. The MC1 megachunk was divided into two to facilitate experiments. **(E)** Assembly of all megachunks on the host chromosome. The unusual *attB-attL* recombination was observed. **(F)** Cre-*loxP* excision of the kernel genome from the host genome chromosome. The small arrows in the genome map indicate the replication fork direction at the strong *ter-*Tus complexes. **(G)** Agarose gel electrophoresis analysis of enzymatically supercoiled chromosomes detected the full-length kernel genome chromosome. BG8, a bipartite-genome strain of *E. coli*, was used for the positive control and the size marker.

### In vitro assembly of the kernel genome components

With this blueprint, we assembled and cloned a complete set of DNA megachunks. Traditionally, Gibson assembly and *E. coli* cloning are used for assembling 10~100 kb DNA chunks, while larger chunks are assembled and cloned by transformation-associated recombination (TAR) using yeast cells (*8, 9, 21, 42*). We used the RA-RCR method (*31, 38*) for seamless cloning and stick to *E. coli* cloning (Figs. 2B, S1). Although the RA-RCR kit was commercially discontinued (OriCiro Genomics, Inc.), we have lab-made reagents. The RA step assembles a circular DNA from linear double-stranded DNA fragments and a linearized BAC vector containing the *E. coli* origin of replication (*oriC*), and the RCR step exclusively amplifies *oriC*-containing circular DNA molecules in an exponential manner (*38*). New chunks are always flanked by novel AvrII sites on the BAC vector and can be excised using AvrII (Figs. 2B, S1). We established a facile method for the purification of AvrII-excised linear DNA fragments as large as 200 kb: Enzymes in reaction mixtures are precipitated with 2-2.5 M ammonium acetate by gentle vortexing and incubation on ice, and after centrifugation the supernatant was ethanol precipitated using sodium acetate and coprecipitant. The proper choice of plasmid miniprep kits greatly facilitated experiments: QIAprep spin columns for small plasmids (up to 20 kb), MonoFas spin columns for large plasmids (up to 200 kb), and NucleoBond PC 20 columns for extra-large plasmids (up to 500 kb). To facilitate >300-kb chunk assembly, we newly developed Tus-masking assembly (Fig. S2). One of the chunk-flanking AvrII sites is overlapped by a *terB* variant sequence (*43*) so that it can be masked by Tus protein during AvrII digestion. Instead, the *oriC* cassette has two AvrII sites to serve as the alternative recombination site. Tus-masking assembly produced a new chunk DNA excisable by AvrII in the absence of Tus.

By repeating the cloning step, we prepared six megachunks from about 360 PCR fragments (Table S1 and Fig. 2C). The chromosomal regions containing the origin (*ori*) and the terminus (*dif*) were fused as a single chromosome with no AvrII site, while the other megachunks ranging from 206 to 423 kb in size are excisable using AvrII (Fig. 2C). The largest chunks are as large as the largest chunk (310 kb) cloned by Gibson assembly (*17, 42*). Although we tried to clone even larger megachunks, we suffered from troubles such as inefficiency, instability, and probably improper joints (data not shown). Rather, we moved to in vivo assembly because we know the portable chromosome format (PCF) according to which a large chromosome can be genetically stabilized in *E. coli* (*31*).

### In vivo assembly of the prototype kernel genome

The *ori-dif* chromosome serves as the core for the integration of the five megachunks and has several devices as follows (Fig. 2D). The wildtype or variant *attB* sites (*attB* or *attBv*) are used for the HK022 integrase-mediated integration of a circular DNA containing a wildtype or variant *attP* site (*attP* or *attPv*), respectively (*26*). The original *oriC* sequence in the *ori* region was replaced by the *ori2* locus of the *Vibrio tubiashii* secondary chromosome to keep the chromosome copy number as low as possible (*26, 44*). An ectopic copy of the *lapC* (*yejM*) gene (*45*) was cloned near the *lapAB* operon, which prevented mutation accumulation in this operon. Preliminary experiments confirmed that the MC2 and MC4 megachunks were integrated into the *ori-dif* core via *attB-attP* or *attBv-attPv* recombination, respectively (Fig. S3). These chromosomes as large as 500 kb were successfully purified and analyzed (Fig. S3). This MC2-*ori-dif* chromosome was used for further experiments (Fig. 2D).

To greatly boost the in vivo assembly of megachunks, we have improved the protocols while actually building larger chromosomes (Fig. 2D). We decided to employ the *oriT*-POP cloning method (*26, 31*) used for the conjugal transfer of a genomic region flanked by two *oriT* sequences. In this study, we optimized this method for plasmid payload delivery (Fig. S4). Each megachunk payload was cloned on an *ori2* BAC vector and flanked by two *oriT* sequences using lambda Red recombination (*46*) (Fig. S4). Instead of the full-length MC1 on an F-plasmid BAC vector, we used the first and second halves (MC1a and MC1b, respectively) separately cloned on the *ori2* vector (Fig. 2D). During the conjugation, the recipient cells are forced to maintain its *ori2* plasmid by antibiotic selection to avoid its replacement by the *ori2* plasmids coming from the donor cells (Fig. S4). Hence, only the *oriT*-flanked megachunk region is accepted by the recipient cell, circularized at *oriT*, and integrated into the recipient’s *ori2* plasmid by the *attB-attP* recombination.

Via plasmid payload delivery, the MC3 megachunk was fused with the MC2-*ori-dif* to make a 0.8-Mb chromosome (Chr^0.8-Mb^) that almost corresponds to the *dif* side of the genome between the ribosomal *rrnGH* operons (*47*). After regeneration (*48*) of the *attB* and *attBv* sequences (Fig. S5), the MC4 and MC1b megachunks were fused with Chr^0.8-Mb^ to make a 1.3-Mb chromosome (Chr^1.3-Mb^). Surprisingly, the NGS analysis of this Chr^1.3-Mb^ revealed its integration into the host chromosome at the genomic *attB* site (Fig. 2E). Besides, errors were found in two non-essential genes; a transposon insertion in the *sbcD* gene and an N-terminal short deletion in the *nuoF* gene (ΔLeu13-Ala41). Then, the integrated 1.3 Mb region was flanked by *loxP* sites to facilitate its excision by Cre (Fig. 2E). The MC1a and MC5 megachunks were integrated to assemble a whole kernel genome (Chr^1.7-Mb^) inside the host chromosome. The Chr^1.7-Mb^ was excised via Cre-*loxP* recombination and transferred to another *E. coli* strain via conjugation (Fig. 2F). To directly check the chromosome size, Chr^1.7-Mb^ was extracted, enzymatically supercoiled by SCR (*31*), and analyzed by canonical agarose gel electrophoresis (Fig. 2G). Fortunately, the Chr^1.7-Mb^ chromosome is no more unstable in cells, possibly owing to balanced gene dosage. The genomic *attB* integration site for the HK022 phage may have served as a genome-building yard to maintain the unstable half-way genome.

### Implementing genome self-digestion in E. coli

We established a host *E. coli* strain that can digest its own genome by the simultaneous multiple-site expression of *avrII* genes upon addition of inducers Crystal Violet (CV) and anhydrotetracycline (aTc) (Fig. 3A). We named this system ‘resident (evil) restriction enzyme (RERE)’ because AvrII is an active restriction enzyme whose leaky expression in cells results in death (18 AvrII sites in the *E. coli* HST08 genome). While the PJExD promoter (*49*) (for CV) and the *tet* promoter of pJKR-H-TetR (*50*) (for aTc) are known as the least leaky promotors for the tight regulation of I-CeuI in *E. coli* (*34*), these promoters were further tuned up for the *avrII* genes (Fig. 3A). We carefully chose their insertion sites on the genome to avoid leaky transcription from outside. Hence, the *avrII* expression cassettes replaced the promoter region and the start codons of oppositely transcribed gene pairs or operons (Fig. 3A). The use of the two novel promoters (PJExD^TT^ and PTetOx2-16) and six genomic safe harbors allowed us to clone three copies of CV-inducible *avrII* genes and three copies of aTc-inducible *avrII* genes in a single strain to establish RERE6 (Fig. 3A). After induction, RERE6 cells became anucleate within 1 hour (Fig. 3B). As expected, the RERE system is practically escape-free in laboratory scale experiments. We did not observe the propagation of escape mutants when we cultured 3.5 x 10^11^ cells for three days in 5.25 L of LB liquid medium containing both CV (1 µM) and aTc (0.1 µ M) (Fig. 3C). After 75 h of induction, the optical density of the medium was only marginally increased, and anucleate cells were observed (Fig. 3C). This 5.25 L culture was estimated to contain at least 1.2 x 10^5^ dormant cells (*51*) that can form colonies on inducer-free LB agar plates. Dormant cells may be negligible as long as the growth medium is supplemented with the inducers.

**Figure 3.**
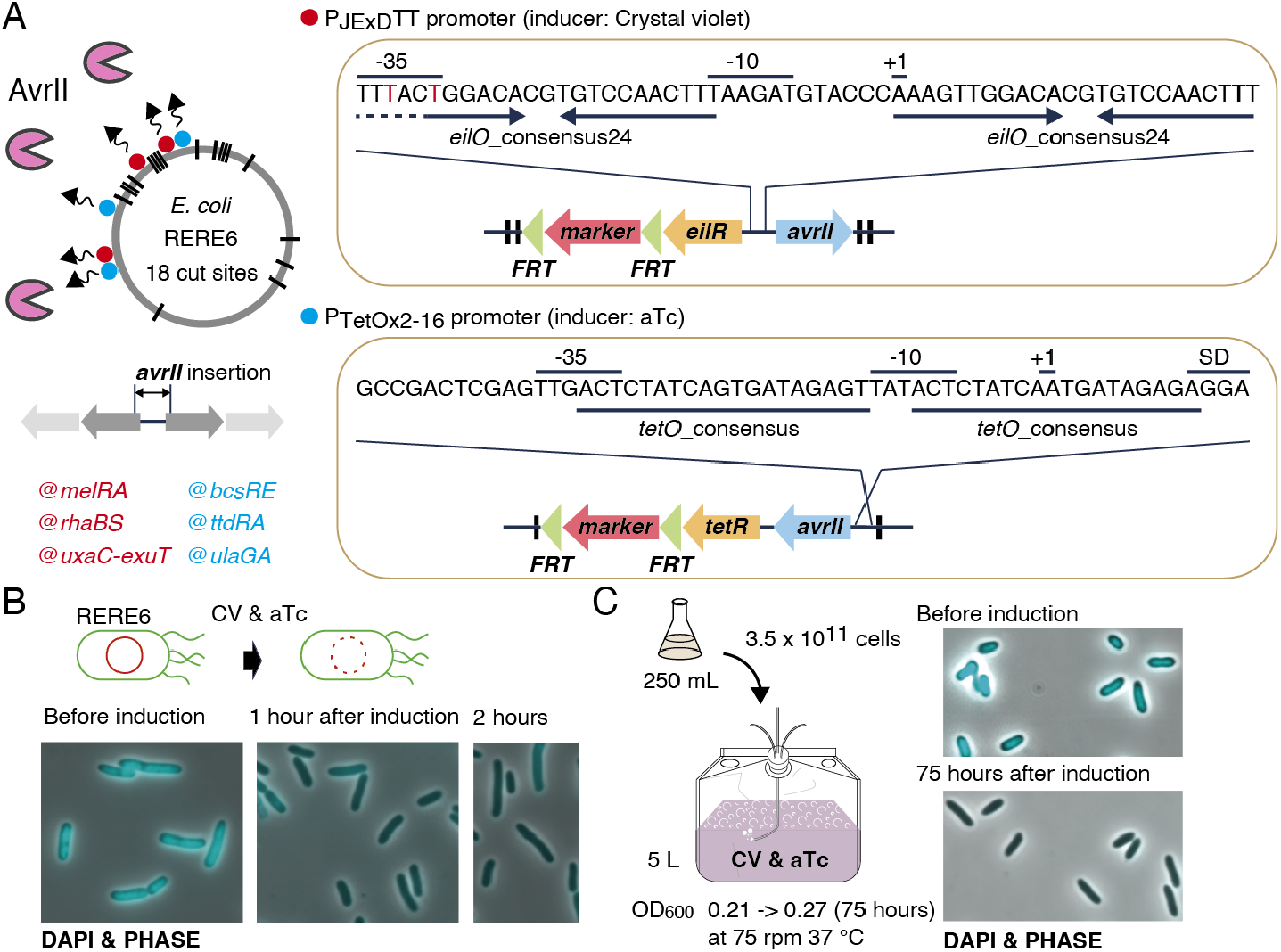
Preparing the host *E. coli* strain installed with the self-digestion system of its own genome. **(A)** *E. coli* RERE6 strain was developed from the HST08 strain by the six-site genomic insertion of the expression cassettes of the *avrII* gene under the control of the EilR and TetR repressors. The selection marker genes flanked with the *FRT* sites can be removed by flippase. **(B)** Chromosome digestion after the AvrII expression induction in RERE6. Cells were no more stainable using DAPI after 1 or 2 hours of the induction. **(C)** Large scale culture experiment of RERE6 cells in the presence of the inducers of the AvrII expression. Propagation of escape mutants was not observed. Anucleate cells were observed after 75 hours of culturing.

### Run and debug the kernel genome

*E. coli* RERE6 cells transformed with the prototype kernel genome were used for its run test and debugging. First, a preliminary run test was performed (Fig. 4A). To facilitate the detection of cells lacking the host chromosome, the mLychee gene (*52*) is cloned in the kernel genome under control of the CV-inducible promoter. Thus, the expression of mLychee implies the simultaneous expression of AvrII eliminating the host chromosome. Weak fluorescent cells were observed after overnight incubation in the presence of CV and aTc in a rich growth medium consisting of LB, M9, glucose, glycerol, and MgSO_4_ (Fig. 4A). However, no colony formation was observed, indicating fatal bugs in the prototype kernel genome.

**Figure 4.**
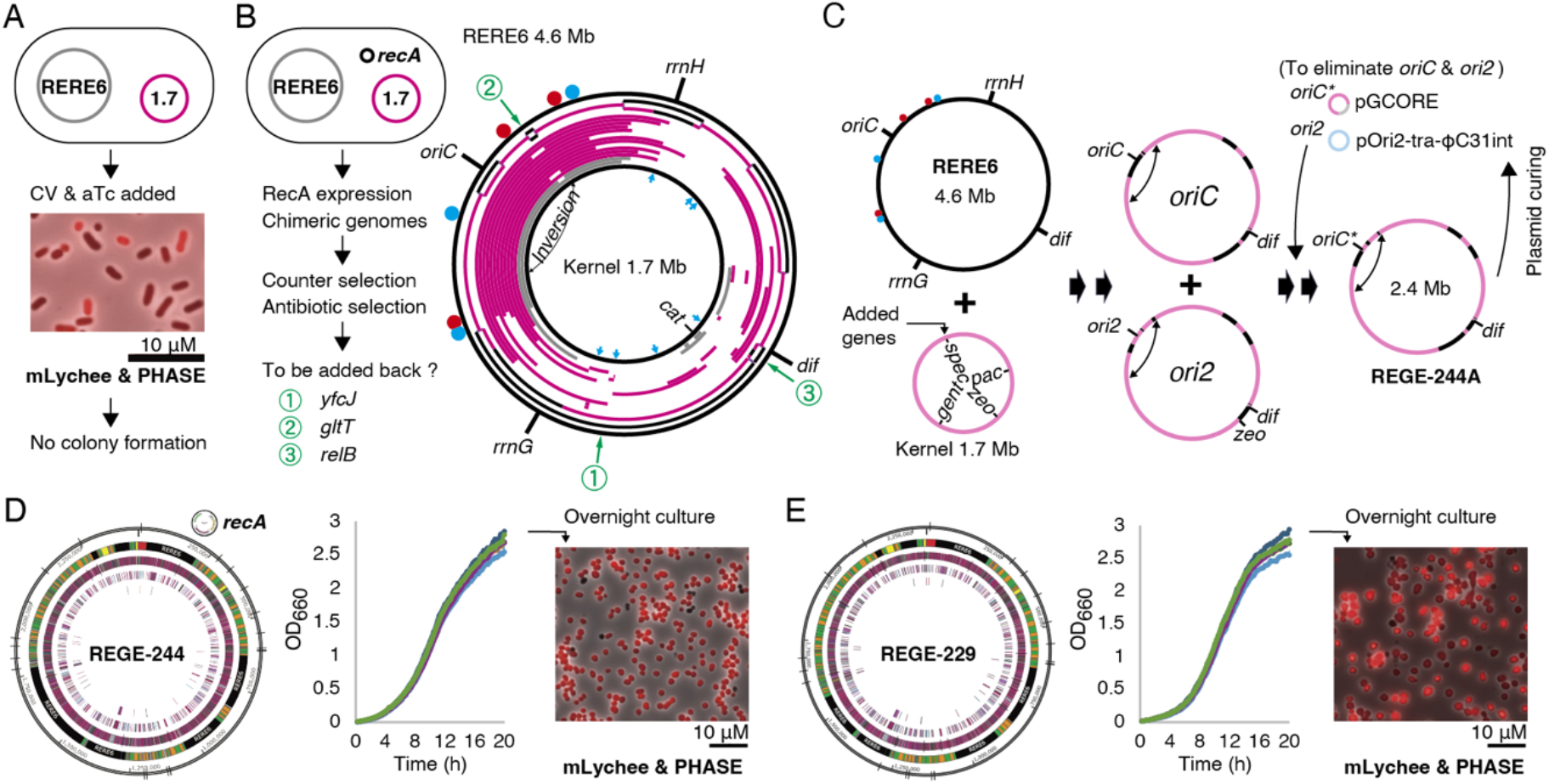
The runtime environment to run and debug the kernel genome. **(A)** RERE6 cells harboring the kernel genome was subjected to a run test. After the induction of AvrII and mLychee from the host genome and the kernel genome, respectively, a portion of living cells were expressing mLychee (red). No colony formation was observed, indicating fatal bugs in the kernel genome. **(B)** RecA-mediated debugging generated chimeras of the RERE6 genome and the kernel genome and identified a few candidate genes to be added back. For each chimera, the inferred kernel-derived regions are mapped on the RERE6 genome. The large inversion between the *<u>rrnDE</u>* operons in the kernel genome derives from the wildtype W3110 strain. Gray lines indicate the original kernel genome lacking the mLychee gene, while red lines indicate mLychee. The outermost chimera genome may be diploid and mosaic, and black lines indicate the RERE6 genome. Circles indicate the six *avrII* genes. Blue arrows indicate the positions of a kanamycin resistance gene inserted into the kernel genome. **(C)** Chimera generation using a modified kernel genome supplemented with several genes and markers. The chimeric chromosomes lack the six *avrII* genes and retain several AvrII sites. The polyploidy of the obtained strain was resolved by forcing the fusion of the multiple chromosomes via transformation with two small chromosomes. pGCORE has a kernel *ori* region containing the *oriC*\* variant and lacks the AvrII sites. pOri2-tra-ΦC31int is a self-transmissible conjugal plasmid expressing ΦC31 integrase under control of the PJExD^TT^ promoter. The *oriC* locus derived from RERE6 is flanked by the ΦC31 *attB* and *attP* sequences and excisable by the integrase. (D) Characterization of the REGE-244A strain. The genome map shows the remaining RERE6 regions in black. The doubling time is 116 (±3) min. Cells in the overnight cultures are mostly oval and red. (E) Characterization of the REGE-229 strain developed by the piecemeal swapping of the REGE-244 genome by the kernel genome via conjugal recombination. The doubling time is 108 (±2) min.

In the presence of RecA, chimeras of the host and kernel genomes were generated and selected by a counter selection using CV and aTc and by a positive selection using antibiotics (Fig. 4B). We analyzed 15 strains by NGS. These chimeras totally lack the host-genome regions containing the six *avrII* genes and have the kernel-genome regions containing the selection markers. The whole RERE6 genome region was separately swapped by the kernel genome in these chimeras. A few loci of the host genome tend to remain intact, and their knockout strains were obtained by isolating a suppressor strain or by enhancing the homologous recombination with ultraviolet light. Careful inspection identified a few candidate genes that should be added back to the kernel genome. The *yfcJ* gene was recently reported as a conditional essential gene for the *bo*_3_ quinol oxidase (*53*). The wildtype *gltT* (tRNA^Glu^) gene may be better than the ΔAvrII variant. The RelB antitoxin gene might be required until the clearance of the RelE toxin. These and a few other candidate genes and supporting genes were cloned to patch the kernel genome via recombinase-mediated cassette exchange (RMCE) (*54*) using a temperature-sensitive mini-F vector (*55*). Despite efforts, we did not obtain chimeric strains controlled solely by an *ori2*-chromosome smaller than 2.5 Mb, close to the size limit of minimal *E. coli* genomes reported (*56, 57*), indicating fatal errors in our genome design or experiments.

We updated our strategy and protocols to enhance chimera generation. The new growth medium is LB (ΔNaCl), M9, glucose, glycerol, MgSO_4_, and glutathione (final 2.5 mg/mL) (*58*). The kernel genome is marked with eight positive selection markers and negative selection markers (two I-SceI sites) (*59*). The AvrII expression was performed in a two-stage manner: The first induction with 1/100 inducers expresses small amount of AvrII for enhancing recombination, while the second induction should eliminate the *avrII* gene-harboring cells. An interesting chimera strain was obtained (Fig. 4C). The strain may harbor a few copies of *oriC*-chromosomes and a single-copy *ori2*-chromosome both of which are mostly derived from the kernel genome (Fig. 4C). This may indicate that the cellular DNA density or the chromosome length does matter, in addition to the individual essential genes and loci. These chromosomes were forcedly fused with another *ori* plasmid pGCORE (*31*) as a single chromosome by disrupting the *oriC* and *ori2* origins via the switcher technology (*60*) and by exploiting the *ori2* incompatibility, respectively (Fig. 4C). The obtained chromosome is 2.44 Mb and powered by the watermarked *oriC* (*oriC*\*) of pGCORE (*31*). This <u>re</u>duced-<u>ge</u>nome strain controlled by the <u>2.44</u> Mb genome with the *rec<u>A</u>*-expressing plasmid is named REGE-244A (Fig. 4D). REGE-244A cells grow slowly but steadily and were oval-shaped rather than rod-like (Fig. 4D).

Lastly, the conjugal recombination of REGE-244 was performed using a kernel genome for further genome trimming (Fig. S6). First, a small region including the I-SceI restriction site in the REGE-244 genome was trimmed by expressing I-SceI and RecA to develop feeble strains (*59*). Meanwhile, the kernel genome in RERE6 was marked with a kanamycin gene at the *argS* gene. Via conjugation, the ΔI-SceI and *argS* region of the feeble strains was replaced by the kernel genome to develop REGE-229 (Figs. 4E, S6). The 2.29 Mb genome is less than half in size compared to the wildtype genome (4.64 Mb) and the smallest ever reported. Nevertheless, REGE-229 grows vigorously (Fig. 4E). REGE-229 can be named as *E. coli* 0.5.

## Discussion

In this study, we developed a genome synthesis platform while actually assembling the half-sized genome. Most importantly, we achieved the direct construction and debugging of a genome-sized chromosome of *E. coli* in *E. coli*. Such a direct experiment has long been limited to yeast. In addition, *E. coli* is able to directly transfer a whole chromosome via conjugation. Thus, the platform may be applicable to constructing a heterologous genome. Furthermore, the platform is compatible with other modern technologies employing yeast or CRISPR/Cas. For example, the half-sized genome could be used as the template or reference for constructing a synthetic artificial genome of *E. coli*. On the other hand, deep inspection of *E. coli* 0.5 would improve our understanding of how *E. coli* lives.

Our platform may have a high affinity with the generative biology field. Our agile tools facilitate rapid prototyping and massive parallel development. More importantly, these tools are designed as user-friendly especially for non-professional biologists and students. Users should not be annoyed by false-positives and mutants escaping the restriction. Genetic systems should be controlled in a deterministic way in a “digital” style. For example, the RERE system is practically escape-free due to the big hurdle to the simultaneous inactivation of the six *avrII* genes. The payload delivery system exploits the strict incompatibility of the *ori2* origin. Very recently, a robust zombie-cell protocol was established for mycoplasmas (*16*). As for *E. coli*, we propose the restriction enzyme-mediated zombification of *E. coli* (REZE) to leave a guest genome intact.

Enfleshing a guest genome is a promising approach for designer genome synthesis. However, the present RTE was limited to chimera generation. Most of the chimeras have emerged through the piecemeal replacement of the host genome by kernel genome fragments. It is suggested that the six *avrII* cassettes of the host genome must have been eliminated prior to the AvrII induction and that toxin-antitoxin loci on the host genome may have forbidden their removal. To partially overcome this limitation, we performed the two-step AvrII induction and added an antitoxin gene. The establishment or use of a toxin-free host strain may be a good solution. The RTE should be updated to enable the patching of the kernel genome with wildtype genome fragments. A customized genome library of *E. coli* may be very useful for this purpose.

The IDE can be updated. As a matter of fact, it took a few years for the construction of the kernel genome. But we expect that a new version could be assembled within a year. A new design principle is to avoid the unexpected emergence of promoter-like sequences at the artificial junctions. In particular, TTGACA, the consensus −35 box, may occur easily. For PCR, the Q5 DNA polymerase should be used to reduce mutations. For constructing middle-sized DNA chunks (40–50 kb), we recommend RA-LAMBDA-INN and iPac (*61, 62*). DNA chunks are assembled in vitro, size-selected on the in vitro packaging into phage particles, and cloneable as a plasmid in *E. coli* by transfection. In principles, only colonies harboring a plasmid of 38-52 kb in size are obtained (see Materials and Methods) (*61*). Megachunks (200–300 kb) can be assembled by Tus-RA-RCR using AvrII-excised DNA chunks and the *ori2* vector. The *ori2* vector is an all-purpose vector and functions even in the genome-integrated state at the genomic *attB* site. A drawback of the *ori2* vector is that one of the artificial *parS2* clusters responsible for plasmid partitioning (*26*) inactivates adjacent regions possibly by forming a heterochromatin-like domain. Thus, we used a very strong promoter for marker genes or inserted a 10-kb spacer between the *parS2* cluster and the selection marker. As for λ Red recombination, it is possible that RecB becomes quasi-essential in *E. coli* harboring a guest genome. Thus, the leaky expression of the RecB-inhibitor GamS from the helper plasmid should be as low as possible.

*E. coli* 0.5 is a proof of concept, not our final goal, because 38.6% of the genome still derives from the wildtype strain. We expect half of the wildtype regions non-essential. Unlike the chemically synthesized minimal genome of JCVI-syn3.0 (*9*), the kernel genome was empirically designed by a human expert, assembled from PCR fragments, and unfortunately incomplete. The next task will be the development of a standalone 2.0 Mb genome via massive patching or trimming to obtain data for AI education, in order to complete the kernel genome. We expect that cells controlled by a small genome would serve as a good host strain to boot up a designer genome owing to the reduced genome-to-genome interference. Developing a bipartite-genome strain may facilitate genomic plasmid transformation (GPT) via electroporation (*30, 31*). In summary, genome size reduction must accelerate the generation and implementation of a genome instance (GIGI).

## Materials and Methods

### General

In general, most of the experiments were performed as reported in our previous works (*26, 30, 31, 38*). The *E. coli* genome was engineered by using λ Red recombination. Plasmids were constructed by using In-Fusion (Takara Bio/Clontech) or RA-RCR (Kurata *et al*., unpublished). Several conjugation helper plasmids were constructed by RA and LAMBDA-INN (NIPPON GENE) in a similar manner to iPac (in vitro Packaging-assisted DNA assembly) (*61, 62*) in order to avoid the PCR amplification of long GC-rich DNA fragments. The transformation and electroporation of *E. coli* cells were performed as reported. *E. coli* HST08/Stellar competent cells (Takara Bio/Clontech) were used. Small plasmids were purified by using QIAprep spin columns (QIAGEN). Large plasmids were purified by using MonoFas spin columns (ANIMOS). Small chromosomes were purified by using NucleoBond PC 20 columns (MACHEREY-NAGEL). Large chromosomes were repaired and supercoiled by SCR and analyzed by agarose gel electrophoresis (*31*). PCR was performed using Q5 (NEB) and sometimes KOD ONE (TOYOBO). Colony PCR was performed using GoTaq (Promega) and sometimes KOD ONE. PCR fragments were purified using NucleoSpin Gel and PCR Clean-up (MACHEREY-NAGEL). Long linear DNA fragments were purified by ammonium acetate precipitation followed by ethanol precipitation using Ethachinmate (NIPPON GENE). Agarose gel electrophoresis was performed as reported. NGS was performed as reported by using iSeq (Illumina) and NEBNext Ultra II FS DNA Library Prep with Multiplex Oligo for Illumina (NEB). Oligo DNAs and small DNA cassettes were purchased from Eurofins. The computer programs are NIH ImageJ, Excel, Photoshop, Illustrator, PyMOL, SnapGene, Geneious Prime. The online servers are PEC (https://shigen.nig.ac.jp/ecoli/pec_w3110/), EcoCyc (https://ecocyc.org/), DFAST (https://dfast.ddbj.nig.ac.jp/), and NEB Tm calculator (https://tmcalculator.neb.com/).

### Plasmids and *E. coli* strains

The making processes were omitted. Please see the plasmid maps and the strain list. pDSG288 and pDSG290 were gifts from Ingmar Riedel-Kruse (Addgene plasmid # 115607 and # 115609, respectively). pJKR-H-tetR was a gift from George Church (Addgene plasmid # 62561). pBAD-I-SceI was a gift from Nikolaus Osterrieder (Addgene plasmid # 60960). The HK022 *xis-int* operon, the *avrII* gene, the PJExD promoter (*49*), and the mLychee gene (*52*) were commercially synthesized. The R6K *ori* and pRK2013 as well as DH5α λpir were obtained from pUTmini-Tn5 Cm Kit (KT-3228, Biomedal). P1*vir* phage was a gift from Yasuhiko Sekine. The temperature-sensitive mutation of mini-F was identified from MG1655 FRT2-1 No.1 recAD:Tc (ME4875 in NBRP). *E. coli* SQ171/pCSacB/ptRNA100 was a gift from Alexander Mankin (Addgene plasmid # 155206). *E. coli* W3110S was obtained from NBRP. The growth curves of *E. coli* were measured by using a compact rocking incubator TVS062CA (Advantec) (*30*). The recipe of the special growth medium: LB (No NaCl) and M9 salts mixed and autoclaved together were supplemented with glucose (final 0.4% w/v), glycerol (final 0.4% v/v), Mg_2_SO_4_ (final 2 mM), and glutathione (reduced form, final 2.5 mg/mL). Glutathione and Crystal violet were from FUJIFILM/Wako. Anhydrotetracycline were from Takara Bio/Clontech. Avoid use of Crystal violet in reusable tubes and flasks.

### Gadgets

For UV-irradiation, we used the UV-C lamp of a clean bench (AS ONE CT-600UVAX) for 5 to 30 seconds. For detecting green/red fluorescent colonies, we used a handheld LED light LED-EXHD (OptoCode).

## Acknowledgments

We thank the technical staff in our laboratory (Rikkyo University): Kanako Saga for assisting with the DNA assembly and Emi Imatani and Kayoko Yamada for assisting with reagent preparation. We also thank Koichi Yano for performing the SCR experiment and Nanako Sakaguchi for analyzing the REGE-244A strain.

## Author contributions

T.M. and M.S. designed and steered the project. M.S. conceived of the restriction enzyme-mediated zombification of *E. coli* and the cell-free genome assembly. T.M. invented all the new methods. All the DNA sequences were designed by MUK-AI (https://orcid.org/0000-0001-8618-598X). T.M., A.O, and E.H. performed experiments and analyzed data. T.M., K.S., and K.Y. developed and analyzed the RERE6 strain. T.M. wrote the manuscript.

## Data availability

Data are available from Dryad (DOI: 10.5061/dryad.r2280gbtm).

## Supplementary data

Table S1. List of PCR chunks.

Table S2. List of *E. coli* strains.

Table S3. List of plasmids and chromosomes.

Table S4. List of DNA cassettes.

Table S5. List of NGS data.

Figure S1. AvrII-RA-RCR method.

Figure S2. Tus-masking assembly method.

Figure S3. Megachunk payload fusion with the *ori-dif* core using two integrase systems.

Figure S4. Plasmid payload delivery.

Figure S5. Setup for *oriT*-POP cloning via λ Red recombination.

Figure S6. Mapping the NGS reads of REGE-229 to the REGE-244A map.

## References

1. Nguyen, E., Poli, M., Durrant, M.G., Kang, B., Katrekar, D., Li, D.B., Bartie, L.J., Thomas, A.W., King, S.H., Brixi, G. et al.. (2024) Sequence modeling and design from molecular to genome scale with Evo. Science, 386, eado9336.

2. King, S.H., Driscoll, C.L., Li, D.B., Guo, D., Merchant, A.T., Brixi, G., Wilkinson, M.E. and Hie, B.L. (2025) Generative design of novel bacteriophages with genome language models. bioRxiv, 2025.2009.2012.675911.

3. Gherman, I.M., Sharma, K., Rees-Garbutt, J., Pang, W., Abdallah, Z.S., Gorochowski, T.E., Grierson, C.S. and Marucci, L. (2025) Accelerated design of Escherichia coli reduced genomes using a whole-cell model and machine learning. Cell Systems, 16.

4. Brixi, G., Durrant, M.G., Ku, J., Naghipourfar, M., Poli, M., Sun, G., Brockman, G., Chang, D., Fanton, A., Gonzalez, G.A. et al.. (2026) Genome modelling and design across all domains of life with Evo 2. Nature, 652, 1349–1361.

5. Koster, C.C., da Costa Oliveira, H., van Beveren, F., Houkes, J., van Hooff, J.J.E., Ettema, T.J.G., van der Oost, J. and Claassens, N.J. (2026) De novo design of synthetic microbial genomes. Nature Reviews Bioengineering, 1–18.

6. Cohen, S.N., Chang, A.C., Boyer, H.W. and Helling, R.B. (1973) Construction of biologically functional bacterial plasmids in vitro. Proc Natl Acad Sci U S A, 70, 3240–3244.

7. Larionov, V., Kouprina, N., Graves, J. and Resnick, M.A. (1996) Highly selective isolation of human DNAs from rodent-human hybrid cells as circular yeast artificial chromosomes by transformation-associated recombination cloning. Proc Natl Acad Sci U S A, 93, 13925–13930.

8. Gibson, D.G., Glass, J.I., Lartigue, C., Noskov, V.N., Chuang, R.Y., Algire, M.A., Benders, G.A., Montague, M.G., Ma, L., Moodie, M.M. et al.. (2010) Creation of a bacterial cell controlled by a chemically synthesized genome. Science, 329, 52–56.

9. Hutchison, C.A., 3rd, Chuang, R.Y., Noskov, V.N., Assad-Garcia, N., Deerinck, T.J., Ellisman, M.H., Gill, J., Kannan, K., Karas, B.J., Ma, L. et al.. (2016) Design and synthesis of a minimal bacterial genome. Science, 351, aad6253.

10. Nucifora, D.P., Saini, T.S. and Karas, B.J. (2026) Direct Genome Transfer from Acholeplasma laidlawii to Yeast. ACS Synthetic Biology, 15, 1353–1360.

11. Taghon, G.J. and Strychalski, E.A. (2023) Rise of synthetic yeast: Charting courses to new applications. Cell Genomics, 3, 100438.

12. Schindler, D., Walker, R.S.K., Jiang, S., Brooks, A.N., Wang, Y., Müller, C.A., Cockram, C., Luo, Y., García, A., Schraivogel, D. et al.. (2023) Design, construction, and functional characterization of a tRNA neochromosome in yeast. Cell, 186, 5237–5253.

13. Zhao, Y., Coelho, C., Hughes, A.L., Lazar-Stefanita, L., Yang, S., Brooks, A.N., Walker, R.S.K., Zhang, W., Lauer, S., Hernandez, C. et al.. (2023) Debugging and consolidating multiple synthetic chromosomes reveals combinatorial genetic interactions. Cell, 186, 5220–5236.

14. Erpf, P.E., Meier, F., Walker, R.S.K., Goold, H.D., Boeke, J.D., Paulsen, I.T. and Pretorius, I.S. (2025) Building synthetic chromosomes one yeast at a time: insights from Sc2.0. Nature Biotechnology, 43, 1911–1918.

15. Lartigue, C., Glass, J.I., Alperovich, N., Pieper, R., Parmar, P.P., Hutchison, C.A., 3rd, Smith, H.O. and Venter, J.C. (2007) Genome transplantation in bacteria: changing one species to another. Science, 317, 632–638.

16. Seidel, Z.P., Assad-Garcia, N., Paralanov, V., Wu, F., Chao, O., Strychalski, E.A., Romantseva, E., Goshia, T., Craig Venter, J. and Glass, J.I. (2026) Selection-free whole genome transplantation revives dead microbes. bioRxiv, 2026.2003.2013.711674.

17. Zhou, J., Wu, R., Xue, X. and Qin, Z. (2016) CasHRA (Cas9-facilitated Homologous Recombination Assembly) method of constructing megabase-sized DNA. Nucleic Acids Research, 44, e124.

18. Venetz, J.E., Del Medico, L., Wölfle, A., Schächle, P., Bucher, Y., Appert, D., Tschan, F., Flores-Tinoco, C.E., van Kooten, M., Guennoun, R. et al.. (2019) Chemical synthesis rewriting of a bacterial genome to achieve design flexibility and biological functionality. Proc Natl Acad Sci U S A, 116, 8070–8079.

19. Wang, T., He, F., He, T., Lin, C., Guan, X., Qin, Z. and Xue, X. (2024) Reconstruction of a robust bacterial replication module. Nucleic Acids Research, 52, 11394–11407.

20. Lajoie, M.J., Rovner, A.J., Goodman, D.B., Aerni, H.R., Haimovich, A.D., Kuznetsov, G., Mercer, J.A., Wang, H.H., Carr, P.A., Mosberg, J.A. et al.. (2013) Genomically recoded organisms expand biological functions. Science, 342, 357–360.

21. Fredens, J., Wang, K., de la Torre, D., Funke, L.F.H., Robertson, W.E., Christova, Y., Chia, T., Schmied, W.H., Dunkelmann, D.L., Beránek, V. et al.. (2019) Total synthesis of Escherichia coli with a recoded genome. Nature, 569, 514–518.

22. Robertson, W.E., Rehm, F.B.H., Spinck, M., Schumann, R.L., Tian, R., Liu, W., Gu, Y., Kleefeldt, A.A., Day, C.F., Liu, K.C. et al.. (2025) Escherichia coli with a 57-codon genetic code. Science, 6771, eady4368.

23. James, J.S., Dai, J., Chew, W.L. and Cai, Y. (2024) The design and engineering of synthetic genomes. Nature Reviews Genetics, 26, 298–319.

24. Zhou, J., Wang, X., Zhou, L., Zhang, H., Ye, S. and Yuan, Y.-J. (2026) Automated linear DNA assembly of A. thaliana’s chloroplast and mitochondrial genome. Nucleic Acids Research, 54, gkag380.

25. Zürcher, J.F., Kleefeldt, A.A., Funke, L.F.H., Birnbaum, J., Fredens, J., Grazioli, S., Liu, K.C., Spinck, M., Petris, G., Murat, P. et al.. (2023) Continuous synthesis of E. coli genome sections and Mb-scale human DNA assembly. Nature, 619, 555–562.

26. Mukai, T., Yoneji, T., Yamada, K., Fujita, H., Nara, S. and Su’etsugu, M. (2020) Overcoming the Challenges of Megabase-Sized Plasmid Construction in Escherichia coli. ACS Synthetic Biology, 9, 1315–1327.

27. Zhong, L., Zhang, Q., Lu, N., Wang, T., Xue, X. and Qin, Z. (2025) The conjugation-associated linear-BAC iterative assembling (CALBIA) method for cloning 2.1-Mb human chromosomal DNAs in bacteria. Cell Research, 35, 309–312.

28. Wang, K., de la Torre, D., Robertson, W.E. and Chin, J.W. (2019) Programmed chromosome fission and fusion enable precise large-scale genome rearrangement and assembly. Science, 365, 922–926.

29. Robertson, W.E., Funke, L.F.H., de la Torre, D., Fredens, J., Wang, K. and Chin, J.W. (2021) Creating custom synthetic genomes in Escherichia coli with REXER and GENESIS. Nature Protocols, 16, 2345–2380.

30. Yoneji, T., Fujita, H., Mukai, T. and Su’etsugu, M. (2021) Grand scale genome manipulation via chromosome swapping in Escherichia coli programmed by three one megabase chromosomes. Nucleic Acids Research, 49, 8407–8418.

31. Fujita, H., Osaku, A., Sakane, Y., Yoshida, K., Yamada, K., Nara, S., Mukai, T. and Su’etsugu, M. (2022) Enzymatic Supercoiling of Bacterial Chromosomes Facilitates Genome Manipulation. ACS Synthetic Biology, 11, 3088–3099.

32. Kobayashi, H. (2018) Regeneration of Escherichia coli from Minicells through Lateral Gene Transfer. Journal of Bacteriology, 200, e00630–17.

33. Wei, E., Jackson-Smith, A. and Endy, D. (2020) Minicells as a potential chassis for engineering lineage-agnostic organisms. bioRxiv, 2020.2007.2031.231670.

34. Lim, B., Yin, Y., Ye, H., Cui, Z., Papachristodoulou, A. and Huang, W.E. (2022) Reprogramming Synthetic Cells for Targeted Cancer Therapy. ACS Synthetic Biology, 11, 1349–1360.

35. Fan, C., Davison, P.A., Habgood, R., Zeng, H., Decker, C.M., Gesell Salazar, M., Lueangwattanapong, K., Townley, H.E., Yang, A., Thompson, I.P. et al.. (2020) Chromosome-free bacterial cells are safe and programmable platforms for synthetic biology. Proc Natl Acad Sci U S A, 117, 6752–6761.

36. Karp, P.D., Paley, S., Caspi, R., Kothari, A., Krummenacker, M., Midford, P.E., Moore, L.R., Subhraveti, P., Gama-Castro, S., Tierrafria, V.H. et al.. The EcoCyc database (2025). EcoSal Plus, 0, eesp–0019-2024.

37. Manzano-Marín, A., Kvist, S. and Oceguera-Figueroa, A. (2023) Evolution of an Alternative Genetic Code in the Providencia Symbiont of the Hematophagous Leech Haementeria acuecueyetzin. Genome Biology and Evolution, 15, evad164.

38. Su’etsugu, M., Takada, H., Katayama, T. and Tsujimoto, H. (2017) Exponential propagation of large circular DNA by reconstitution of a chromosome-replication cycle. Nucleic Acids Research, 45, 11525–11534.

39. Hirokawa, Y., Kawano, H., Tanaka-Masuda, K., Nakamura, N., Nakagawa, A., Ito, M., Mori, H., Oshima, T. and Ogasawara, N. (2013) Genetic manipulations restored the growth fitness of reduced-genome Escherichia coli. J Biosci Bioeng, 116, 52–58.

40. Uhlenbeck, O.C. and Schrader, J.M. (2018) Evolutionary tuning impacts the design of bacterial tRNAs for the incorporation of unnatural amino acids by ribosomes. Curr Opin Chem Biol, 46, 138–145.

41. Quan, S., Skovgaard, O., McLaughlin, R.E., Buurman, E.T. and Squires, C.L. (2015) Markerless Escherichia coli rrn Deletion Strains for Genetic Determination of Ribosomal Binding Sites. G3 (Bethesda), 5, 2555–2557.

42. Gibson, D.G., Young, L., Chuang, R.Y., Venter, J.C., Hutchison, C.A., 3rd and Smith, H.O. (2009) Enzymatic assembly of DNA molecules up to several hundred kilobases. Nature Methods, 6, 343–345.

43. Coskun-Ari, F.F. and Hill, T.M. (1997) Sequence-specific Interactions in the Tus-Ter Complex and the Effect of Base Pair Substitutions on Arrest of DNA Replication in Escherichia coli. Journal of Biological Chemistry, 272, 26448–26456.

44. Messerschmidt, S.J., Schindler, D., Zumkeller, C.M., Kemter, F.S., Schallopp, N. and Waldminghaus, T. (2016) Optimization and Characterization of the Synthetic Secondary Chromosome synVicII in Escherichia coli. Front Bioeng Biotechnol, 4, 96.

45. Klein, G., Wieczorek, A., Szuster, M. and Raina, S. (2021) Checkpoints That Regulate Balanced Biosynthesis of Lipopolysaccharide and Its Essentiality in Escherichia coli. Int J Mol Sci, 23, 189.

46. Thomason, L.C., Costantino, N., Li, X. and Court, D.L. (2023) Recombineering: Genetic Engineering in Escherichia coli Using Homologous Recombination. Curr Protoc, 3, e656.

47. Takada, H., Shimada, T., Dey, D., Quyyum, M.Z., Nakano, M., Ishiguro, A., Yoshida, H., Yamamoto, K., Sen, R. and Ishihama, A. (2016) Differential Regulation of rRNA and tRNA Transcription from the rRNA-tRNA Composite Operon in Escherichia coli. PLoS One, 11, e0163057.

48. Norville, J.E., Gardner, C.L., Aponte, E., Camplisson, C.K., Gonzales, A., Barclay, D.K., Turner, K.A., Longe, V., Mincheva, M., Teramoto, J. et al.. (2016) Assembly of Radically Recoded E. coli Genome Segments. bioRxiv, 070417.

49. Ruegg, T.L., Pereira, J.H., Chen, J.C., DeGiovanni, A., Novichkov, P., Mutalik, V.K., Tomaleri, G.P., Singer, S.W., Hillson, N.J., Simmons, B.A. et al.. (2018) Jungle Express is a versatile repressor system for tight transcriptional control. Nature Communications, 9, 3617.

50. Rogers, J.K., Guzman, C.D., Taylor, N.D., Raman, S., Anderson, K. and Church, G.M. (2015) Synthetic biosensors for precise gene control and real-time monitoring of metabolites. Nucleic Acids Research, 43, 7648–7660.

51. Kim, J.S., Chowdhury, N., Yamasaki, R. and Wood, T.K. (2018) Viable but non-culturable and persistence describe the same bacterial stress state. Environ Microbiol, 20, 2038–2048.

52. Fraikin, N., Couturier, A., Mercier, R. and Lesterlin, C. (2025) A palette of bright and photostable monomeric fluorescent proteins for bacterial time-lapse imaging. Science Advances, 11, eads6201.

53. Khalfaoui-Hassani, B., Blaby-Haas, C.E., Verissimo, A. and Daldal, F. (2023) The Escherichia coli MFS-type transporter genes yhjE, ydiM, and yfcJ are required to produce an active bo3 quinol oxidase. PLOS ONE, 18, e0293015.

54. Urtecho, G., Tripp, A.D., Insigne, K.D., Kim, H. and Kosuri, S. (2019) Systematic Dissection of Sequence Elements Controlling σ70 Promoters Using a Genomically Encoded Multiplexed Reporter Assay in Escherichia coli. Biochemistry, 58, 1539–1551.

55. Kato, J.i. and Hashimoto, M. (2007) Construction of consecutive deletions of the Escherichia coli chromosome. Molecular Systems Biology, 3, 132.

56. Kotaka, Y., Hashimoto, M., Lee, K.-i. and Kato, J.-i. (2023) Mutations identified in engineered Escherichia coli with a reduced genome. Frontiers in Microbiology, 14, 1189877.

57. Kim, K., Choe, D., Cho, S., Palsson, B. and Cho, B.-K. (2024) Reduction-to-synthesis: the dominant approach to genome-scale synthetic biology. Trends in Biotechnology, 42, 1048–1063.

58. Kawai, Y., Mercier, R., Wu Ling J., Domínguez-Cuevas, P., Oshima, T. and Errington, J. (2015) Cell Growth of Wall-Free L-Form Bacteria Is Limited by Oxidative Damage. Current Biology, 25, 1613–1618.

59. Chayot, R., Montagne, B., Mazel, D. and Ricchetti, M. (2010) An end-joining repair mechanism in Escherichia coli. Proc Natl Acad Sci U S A, 107, 2141–2146.

60. Kasari, M., Kasari, V., Kärmas, M. and Jõers, A. (2022) Decoupling Growth and Production by Removing the Origin of Replication from a Bacterial Chromosome. ACS Synthetic Biology, 11, 2610–2622.

61. Nozaki, S. (2022) Rapid and Accurate Assembly of Large DNA Assisted by In Vitro Packaging of Bacteriophage. ACS Synthetic Biology, 11, 4113–4122.

62. Nozaki, S. and Miwa, Y. (2026) PhAGE Enables One-Step Genome Integration of Large DNA Fragments in Escherichia coli. bioRxiv, 2026.2004.2023.720475.

