## Supplementary for "Generating *E. coli* 0.5 controlled by a half-sized genome"

*Co-corresponding authors.

**Supplementary Tables**

Table S1. List of PCR chunks.

Table S2. List of *E. coli* strains.

Table S3. List of plasmids and chromosomes.

Table S4. List of DNA cassettes.

Table S5. List of NGS data.

**Supplementary Figures**

Figure S1. AvrII-RA-RCR method.

Figure S2. Tus-masking assembly method.

Figure S3. Megachunk payload fusion with the *ori-dif* core using two integrase systems.

Figure S4. Plasmid payload delivery.

Figure S5. Setup for *oriT*-POP cloning via λ Red recombination.

Figure S6. Mapping the NGS reads of REGE-229 to the REGE-244A map.

**Table S1. List of PCR chunks.** The template genome is RGF008C or indicated.

| chunk name | size (bp) | origin | megachunk |
|---|---|---|---|
| N-1to3 | 29823 | | ori |
| N-4 edited | 12029 | | MC1a |
| N-5-1 | 2316 | | MC1a |
| N-5-2 | 8087 | | MC1a |
| N-6 | 12388 | | MC1a |
| N-7-1 | 3914 | | MC1a |
| N-7-2 | 1389 | | MC1a |
| N-7-3 | 3139 | | MC1a |
| N-8-1 | 8230 | | MC1a |
| N-8-2 edited | 1309 | | MC1a |
| N-9-1 | 3088 | | MC1a |
| N-9-2 | 1290 | MG1655 | MC1a |
| N-10-1 | 10334 | | MC1a |
| N-10-2 | 2780 | MG1655 | MC1a |
| O-1-1 | 2092 | | MC1a |
| O-1-2 | 5197 | | MC1a |
| O-1-3 | 886 | | MC1a |
| O-1-4 | 1398 | | MC1a |
| O-1-5 | 1898 | | MC1a |
| O-2-1 | 2387 | | MC1a |
| O-2-2 to feoB | 741 | | MC1a |
| O-2-2 from feoB | 9837 | | MC1a |
| O-3-1 | 2183 | | MC1a |
| O-3-2 | 2903 | | MC1a |
| O-4-1 | 7040 | | MC1a |
| O-4-2 | 2107 | | MC1a |
| O-4-3 | 1476 | | MC1a |
| O-5and6 | 13129 | | MC1a |
| O-7 | 14530 | | MC1a |
| O-8 extended | 17087 | W3110S | MC1a |
| O-9to10 | 20430 | | MC1a |
| P-1-1 edited | 2113 | | MC1a |
| P-1-2 | 5194 | | MC1a |
| P-1-3 | 3897 | | MC1a |
| P-2-1 | 1181 | MG1655 | MC1a |
| P-2-2 | 5029 | MG1655 | MC1a |
| P-2-3 | 304 | | MC1a |
| P-2-4 | 2201 | | MC1a |
| P-3-1 edited | 2259 | | MC1b |
| P-3-2 | 2024 | | MC1b |
| P-3-3 | 1092 | | MC1b |
| P-3-4 edited | 182 | | MC1b |
| P-3-5 edited | 4679 | | MC1b |
| P-4to5 edited to mutL | 3366 | MG1655 | MC1b |
| P-4to5 edited from mutL | 12645 | MG1655 | MC1b |
| P-6-1 | 1775 | | MC1b |
| P-6-2 | 804 | | MC1b |
| P-6-3 | 881 | | MC1b |
| P-6-4 | 3085 | | MC1b |
| P-6-5 edited | 1369 | | MC1b |
| P-6-6 | 1558 | | MC1b |
| P-7-1 | 3643 | | MC1b |
| P-7-2 | 2531 | | MC1b |

| | | | |
|---|---|---|---|
| P-7-3 | 1936 | | MC1b |
| P-8-1 | 7693 | | MC1b |
| P-8-2 | 320 | | MC1b |
| P-8-3 edited | 601 | MG1655 | MC1b |
| P-8-4 | 1408 | MG1655 | MC1b |
| P-9-1 to dnaCT | 782 | | MC1b |
| P-9-1 from dnaCT | 667 | | MC1b |
| P-9-2&3 edited | 6042 | | MC1b |
| P-9-4 edited | 523 | | MC1b |
| P-9-5 | 4559 | | MC1b |
| P-10 | 12401 | | MC1b |
| A-1-1 | 7299 | | MC1b |
| A-1-2 | 3420 | | MC1b |
| A-2 | 13450 | | MC1b |
| new A-3to4 | 17348 | | MC1b |
| new A-5to6 | 20663 | | MC1b |
| new A-7 | 14394 | | MC1b |
| new A-8 | 11718 | | MC1b |
| A-9-1 | 5830 | | MC1b |
| A-9-2 | 1489 | | MC1b |
| A-9-3 | 647 | | MC1b |
| A-9-4 | 1934 | | MC1b |
| A-10-1 | 3532 | | MC1b |
| B-1 | 11866 | | MC1b |
| B-2 | 11489 | | MC1b |
| B-3 | 12074 | | MC1b |
| B-4 | 11544 | | MC1b |
| B-5-1 | 666 | | MC2 |
| B-5-2 | 1708 | | MC2 |

| | | | |
|---|---|---|---|
| B-5-3 | 1427 | | MC2 |
| B-5-4 | 3600 | | MC2 |
| B-6 | 10170 | | MC2 |
| B-7-1 | 1122 | | MC2 |
| B-7-2 | 6702 | | MC2 |
| B-7-3 | 6273 | | MC2 |
| B-8 | 10920 | | MC2 |
| B-9 | 9940 | | MC2 |
| B-10 | 8974 | | MC2 |
| C-1-1 | 2007 | | MC2 |
| C-1-2 | 1237 | | MC2 |
| C-2 | 11823 | | MC2 |
| C-3-1 | 840 | | MC2 |
| C-3-2 | 6803 | | MC2 |
| C-3-3 | 660 | | MC2 |
| C-3-4 | 3517 | | MC2 |
| C-3-5 | 725 | MG1655 | MC2 |
| C-3-6 | 1825 | MG1655 | MC2 |
| C-3-7 | 557 | MG1655 | MC2 |
| new C-4 | 15748 | | MC2 |
| new C-5 | 1121 | MG1655 | MC2 |
| new C-6 | 12226 | | MC2 |
| new C-7 to glnS | 5821 | | MC2 |
| new C-7 from glnS | 8052 | | MC2 |
| C-8-1 | 2281 | MG1655 | MC2 |
| C-8-2 | 887 | MG1655 | MC2 |
| C-9 | 12215 | | MC2 |
| C-10 | 12293 | | MC2 |
| D-1-1 | 2360 | | MC2 |

| | | | |
|---|---|---|---|
| D-1-2 | 1231 | | MC2 |
| D-1-3 | 7835 | | MC2 |
| D-2-1 | 3861 | MG1655 | MC2 |
| D-2-2 | 1141 | MG1655 | MC2 |
| D-2-3 | 996 | | MC2 |
| D-2-4 | 1165 | | MC2 |
| D-2-5 | 1533 | MG1655 | MC2 |
| D-2-6 | 825 | | MC2 |
| D-2-7 | 496 | | MC2 |
| D-3-1 | 2029 | MG1655 | MC2 |
| D-3-2 | 1948 | | MC2 |
| D-3-3 | 6962 | | MC2 |
| D-4 | 11630 | | MC2 |
| D-5to6 | 19460 | | MC2 |
| D-7to8 | 19034 | | MC2 |
| D-9-1 | 1840 | | MC2 |
| D-9-2 | 4263 | | MC2 |
| D-9-3 | 6606 | | MC2 |
| D-10 | 3985 | | MC2 |
| E-1-1 | 475 | | MC2 |
| E-1-2 | 237 | | MC2 |
| E-1-3 | 3460 | | MC2 |
| E-2-1 | 2939 | | MC2 |
| E-2-2 | 716 | | MC2 |
| E-2-3 | 7297 | | MC2 |
| E-3 | 10212 | | MC2 |
| E-4-1 | 1927 | | MC2 |
| E-4-2 | 5642 | | MC2 |
| E-4-3 | 3685 | | MC2 |

| | | | |
|---|---|---|---|
| E-5-1 | 5254 | | MC2 |
| E-6-1 | 9509 | | MC2 |
| E-6-2 | 2434 | | dif |
| ycgJ | 566 | | dif |
| E-7-1 | 3323 | | dif |
| E-7-2 | 335 | | dif |
| E-7-3 | 1103 | | dif |
| E-7-4 | 1629 | | dif |
| E-7-5 | 2165 | | dif |
| ychHM | 2143 | | dif |
| E-8-1 | 8320 | | dif |
| E-8-2 | 1784 | | dif |
| E-9 added | 9873 | | dif |
| E-10-1 | 2785 | | dif |
| E-10-2 | 7799 | | dif |
| F-1-1 full | 5152 | | dif |
| F-1-2-2-1 | 12828 | | dif |
| yajKLM | 3242 | | dif |
| F-1-2-2-1 | 60 | | dif |
| F-2full | 14615 | | dif |
| F-3-1 | 1803 | MG1655 | dif |
| F-3-2 | 2360 | | dif |
| F-3-3 | 538 | MG1655 | dif |
| F-3-4 | 1033 | MG1655 | dif |
| F-3-5 | 3927 | MG1655 | dif |
| F-4-1 | 3547 | MG1655 | dif |
| F-4-2 | 795 | MG1655 | dif |
| F-4-3 | 2077 | MG1655 | dif |
| F-4-4 | 3335 | | dif |

| | | | |
|---|---|---|---|
| F-5-1 | 5694 | | dif |
| F-5-2 | 278 | | dif |
| F-5-3 | 2384 | MG1655 | dif |
| F-5-4 | 360 | | dif |
| F-5-5 | 1398 | MG1655 | dif |
| F-6-1 | 2334 | | dif |
| F-6-2 | 3188 | | dif |
| F-6-3 | 4477 | | dif |
| F-7-1 | 1329 | | dif |
| F-7-2 | 7736 | | dif |
| F-8-1full | 11076 | | dif |
| F-8-3 | 1298 | | dif |
| F-9-1 | 2546 | | dif |
| F-9-2 | 1674 | | dif |
| F-10-1 | 2682 | | dif |
| F-10-2 | 448 | | dif |
| F-10-3 | 9865 | | dif |
| ratAB | 757 | | oridif |
| yadGH | 1713 | | oridif |
| S3added | 730 | | MC3 |
| G-1-1 | 946 | | MC3 |
| G-1-2 | 3654 | | MC3 |
| G-2 | 10550 | | MC3 |
| PthrS-TydiY | 2661 | | MC3 |
| G-3-1 | 5644 | | MC3 |
| G-3-2 | 4110 | | MC3 |
| G-3-3 | 1048 | | MC3 |
| G-4-1 | 5482 | | MC3 |
| G-4-2 | 4526 | MG1655 | MC3 |

| | | | |
|---|---|---|---|
| G-5-1 | 4341 | | MC3 |
| G-5-2 | 731 | MG1655 | MC3 |
| G-6-1 | 8231 | | MC3 |
| G-6-2 | 1032 | | MC3 |
| G-7 | 11895 | | MC3 |
| yebY | 1505 | | MC3 |
| G-8-1 | 459 | | MC3 |
| G-8-2 | 675 | | MC3 |
| G-8-3 | 5657 | | MC3 |
| G-8-4 | 4485 | | MC3 |
| G-9-1 | 6669 | | MC3 |
| G-9-2 | 1962 | | MC3 |
| G-9-3 | 442 | | MC3 |
| G-9-4 | 1996 | | MC3 |
| G-10-1 | 1209 | | MC3 |
| G-10-2 | 2689 | MG1655 | MC3 |
| G-10-3 | 840 | MG1655 | MC3 |
| G-10-4 | 1168 | MG1655 | MC3 |
| tcyJ | 3458 | MG1655 | MC3 |
| yeeX | 601 | MG1655 | MC3 |
| yciU | 422 | MG1655 | MC3 |
| yfbV | 473 | MG1655 | MC3 |
| yecM | 633 | MG1655 | MC3 |
| G-10-4 | 129 | MG1655 | MC3 |
| G-10-5 | 1503 | | MC3 |
| G-10-6 | 285 | | MC3 |
| G-10-7 | 257 | | MC3 |
| G-10-8 | 2654 | | MC3 |
| G-10-9 | 1704 | | MC3 |

| | | | |
|---|---|---|---|
| H-1-1 | 7476 | | MC3 |
| H-1-2 | 3048 | | MC3 |
| H-2-1 | 3800 | | MC3 |
| H-2-2 | 4369 | MG1655 | MC3 |
| H-2-3 | 2114 | MG1655 | MC3 |
| H-3-1 | 1923 | | MC3 |
| H-3-2 | 3524 | | MC3 |
| H-3-3 | 1027 | | MC3 |
| H-3-4 | 1945 | | MC3 |
| H-3-5 | 402 | | MC3 |
| H-3-6 | 1870 | | MC3 |
| H-4-1 | 1032 | | MC3 |
| H-4-2 | 2514 | | MC3 |
| H-4-3 | 1948 | | MC3 |
| H-4-4 | 6621 | | MC3 |
| H-5-1 | 4567 | | MC3 |
| H-5-2 | 3876 | | MC3 |
| H-6-1 | 5719 | | MC3 |
| H-6-2 | 1926 | MG1655 | MC3 |
| H-6-3 | 3121 | MG1655 | MC3 |
| H-7 | 12701 | | MC3 |
| H-8-1 | 2277 | | MC3 |
| H-8-2 | 7675 | | MC3 |
| H-9-1 | 7296 | | MC3 |
| H-9-2 | 4390 | | MC3 |
| H-10-1 | 2920 | | MC3 |
| H-10-2 | 1642 | | MC3 |
| H-10-3 | 1081 | | MC3 |
| H-10-4 | 1566 | | MC3 |

| | | | |
|---|---|---|---|
| nupC | 1516 | | MC3 |
| I-1-1 | 441 | | MC3 |
| cysPUWA | 4151 | | MC3 |
| I-1-2 | 2394 | | MC3 |
| I-1-3 | 9250 | | MC3 |
| I-2-1 | 4957 | | MC3 |
| I-2-2 | 2025 | | MC3 |
| I-2-3 | 5891 | | MC3 |
| I-3 | 9589 | | MC3 |
| yfgO | 1218 | | MC3 |
| I-4-1 | 2995 | | MC3 |
| I-4-2 | 6530 | | MC3 |
| I-4-3 | 773 | | MC3 |
| I-5 | 14225 | | MC3 |
| I-6-1 | 1147 | | MC3 |
| I-6-2 | 7824 | | MC3 |
| I-6-3 | 1742 | | MC3 |
| I-7-1 | 8699 | | MC3 |
| I-7-2 | 1475 | | MC3 |
| I-7-3 | 319 | | MC3 |
| I-8 | 14079 | | MC3 |
| I-9 | 7585 | | MC3 |
| J-1 | 9844 | | MC4 |
| J-2-1 | 3874 | | MC4 |
| J-2-2 | 4100 | | MC4 |
| J-2-3 | 1417 | | MC4 |
| J-3-1 | 8056 | | MC4 |
| J-3-2 | 715 | MG1655 | MC4 |
| J-3-3 | 3575 | | MC4 |

| | | | |
|---|---|---|---|
| J-4 | 10091 | | MC4 |
| J-5 | 5136 | | MC4 |
| J-6-1 | 5257 | | MC4 |
| J-6-2 | 6736 | | MC4 |
| J-7-1 | 3201 | MG1655 | MC4 |
| J-7-2 | 1761 | MG1655 | MC4 |
| J-7-3 | 1970 | MG1655 | MC4 |
| J-7-4 | 4126 | | MC4 |
| J-8-1 | 6782 | | MC4 |
| J-8-2 | 3369 | | MC4 |
| J-9-1 | 1950 | | MC4 |
| J-9-2 | 1905 | | MC4 |
| J-9-3 | 3971 | | MC4 |
| J-9-4 | 2443 | | MC4 |
| J-10 | 9679 | BL21(DE3) | MC4 |
| added to K-1 | 891 | | MC4 |
| K-1-1 edited | 5869 | | MC4 |
| K-1-2 | 2394 | MG1655 | MC4 |
| K-1-3 | 1023 | MG1655 | MC4 |
| K-1-4 | 4020 | | MC4 |
| K-2 edited | 8457 | | MC4 |
| K-3-1 | 4898 | | MC4 |
| K-3-2 | 714 | | MC4 |
| K-3-3 | 3819 | | MC4 |
| K-3-4 | 2419 | | MC4 |
| K-3-5 | 329 | | MC4 |
| K-3-6 | 409 | | MC4 |
| K-4-1 to metC | 2952 | | MC4 |
| K-4-1 from metC | 4147 | | MC4 |

| | | | |
|---|---|---|---|
| K-4-2 edited | 4789 | | MC4 |
| K-5 | 11845 | | MC4 |
| K-6 | 9374 | | MC4 |
| K-7-1 edited | 6375 | | MC4 |
| K-7-2 | 2150 | | MC4 |
| K-7-3 | 1401 | MG1655 | MC4 |
| K-8-1 | 2899 | | MC4 |
| K-8-2 edited | 1135 | MG1655 | MC4 |
| K-8-3 | 4805 | | MC4 |
| K-8-4 | 1422 | | MC4 |
| K-9 edited | 3107 | MG1655 | MC4 |
| K-10 edited | 11171 | | MC4 |
| L-1to2 | 18321 | | MC4 |
| L-3to4 | 20563 | | MC4 |
| satP | 684 | | MC5 |
| panE-yajL | 1486 | | MC5 |
| tamAB | 8563 | MG1655 | MC5 |
| L-5 | 9737 | | MC5 |
| yhdP | 3868 | | MC5 |
| L-6 | 7161 | | MC5 |
| L-7-1 | 6677 | | MC5 |
| L-7-2 | 3059 | | MC5 |
| L-8 to hemE | 1931 | | MC5 |
| L-8 from hemE | 13069 | | MC5 |
| L-9 | 13000 | | MC5 |
| L-10 | 10095 | | MC5 |
| L-11 | 9343 | | MC5 |
| M-1-1 | 2904 | | MC5 |
| Re M-1-2toM-2-3 | 25385 | | MC5 |

| | | | |
|---|---|---|---|
| M-2-4 | 1056 | | MC5 |
| M-3-1 | 1430 | | MC5 |
| M-3-2 | 3036 | | MC5 |
| M-3-3 | 792 | | MC5 |
| M-3-4 | 458 | MG1655 | MC5 |
| M-3-5 edited | 6926 | | MC5 |
| igaA | 2420 | | MC5 |
| slyD | 974 | | MC5 |
| M-3-5 edited | 40 | | MC5 |
| M-4-1 | 5034 | | MC5 |
| M-4-3 edited | 2076 | | MC5 |
| M-5 | 10294 | | MC5 |
| M-6 | 13589 | | MC5 |
| M-7-1 | 4437 | | MC5 |
| M-7-2 | 2940 | | MC5 |
| M-7-3 | 1359 | | MC5 |
| M-8 | 13983 | | MC5 |
| M-9-1 | 690 | | MC5 |
| M-9-2 edited to rho | 1091 | | MC5 |
| M-9-2 edited from rho to M-10 | 16530 | | MC5 |

**Table S2. List of *E. coli* strains.**

| strain number | strain name | description |
|---|---|---|
| RGF008C | RGF008C | DGF-298W Δ*sacB-cat* Δ*terFIJ* Δ*recAX* (ref. Yoneji et al, 2021) |
| GP80 | HST08 ΔattB | Lacking the HK022 *attB* (ref. Mukai et al., 2020) |
| GP105 | W3110S (NIG Wide Deletion Strain Collection from NBRP) | One of the wildtype *E. coli* strains (ref. Hirokawa et al., 2013) |
| GP153 | *E. coli* BG8 | a bipartite-genome strain of *E. coli* (ref. Fujita et al., 2022) |
| GP192 | HST08 ΔattB endA1::tet | conjugation recipient strain |
| GP207 | SQ171/pMW118cat-rrnCΔAvrII/ptRNA100-gltT::zeo | The *rrn* and tRNA plasmids of SQ171(Addgene 155206) were replaced. |
| GP218 | GP192 attB-oriC-attP gyrAwt | The *oriC* is flanked by the φC31 *attBP*. |
| GP270 | RERE6 (GP218 with the 6 *avrII* cassettes) | no antibiotic marker |
| GP279 | RERE6kzs (still having the anitibiotic markers in the aTc-avrII cassetts) | *kan, zeo, spec* |
| XGP_02 | HST08 pOri2-oridif clone: 1B3 | 0.2 Mb |
| XGP_05 | HST08 MC2-oridif pMWcat | 0.5 Mb |
| XGP_08 | HST08 MC2-oridif-MC3-attB gent-gba | 0.8 Mb |
| XGP_10 | HST08 MC2-oridif-MC3-MC4-attB-attBv pMW118 | 1.08 Mb |
| XGP_13p | 1.3 Mb pTra2013 pTet17-HKint clone: 14-24-8 | for NGS |
| XGP_13g | 1.3 Mb in genome pTra2013 hyg-gba clone: i-a | for NGS |
| XGP_17 | 1.7 Mb in genome No. 1-4 | for NGS |
| GP333 | RERE6 pJExDTT-NoeilR-sfGFP-TtufB_2xAvrII | To confer gentamicin resistance. |
| GP344 | GP192 pMW118-g'ba | conjugation recipient strain |
| GP346 | GP192 Kernel pMW118-g'ba | GP192 received the original kernel genome. |
| GP347 | GP333 Kernel | GP333 received the original kernel genome. |
| XGP_8 | chimera_8 | A chimeric strain derived from GP333 |
| XGP_24 | chimera_24 | A chimeric strain derived from GP333 |
| GP349 | GP347 hyg-gba kernel-attPv-zeo-oriT::attBv-mLychee-bla | The original mLychee kernel genome |
| GP353 | GP347 hyg-gba kernel-attPv-zeo-oriT::attBv-mLychee-bla kan@B-4/5 | |
| GP355 | GP347 hyg-gba kernel-attPv-zeo-oriT::attBv-mLychee-bla kan@C-8-1/2 | |

| | | |
|---|---|---|
| GP365 | chimera_C-8-1/2-1 | A chimeric strain derived from GP355 |
| GP366 | chimera_C-8-1/2-2 | A chimeric strain derived from GP355 |
| GP369 | GP347 hyg-gba kernel-attPv-zeo-oriT::attBv-mLychee-bla kan@G/H | |
| GP370 | GP347 hyg-gba kernel-attPv-zeo-oriT::attBv-mLychee-bla kan@I-1-1/cys | |
| GP372 | chimera_G/H-1 | A chimeric strain derived from GP369 |
| GP373 | chimera_G/H-2 | A chimeric strain derived from GP369 |
| GP375 | chimera_B-4/5-e | A chimeric strain derived from GP353 |
| GP377 | chimera_I/Cys-old | A chimeric strain derived from GP370 |
| GP378 | chimera_I/Cys-new-a | A chimeric strain derived from GP370 |
| GP379 | chimera_I/Cys-u | A chimeric strain derived from GP370 |
| XGP_38 (discarded) | GP347 hyg-gba kernel-attPv-zeo-oriT::attBv-mLychee-bla kan@H-7/8 | |
| GP380 | chimera_H-7/8 No. 4 | A chimeric strain derived from XGP_38 |
| GP381 | chimera_H-7/8 No. 7 | A chimeric strain derived from XGP_38 |
| GP387 | GP347 hyg-gba kernel-attPv-zeo-oriT::attBv-mLychee-bla kan@C5/6 | |
| GP388 | GP347 hyg-gba kernel-attPv-zeo-oriT::attBv-mLychee-bla kan@F5/6 | |
| GP389 | chimera_C-5/6 No. 13 | A chimeric strain derived from GP387 |
| GP390 | chimera_F-5/6 No. 22 | A chimeric strain derived from GP388 |
| GP391 | chimera_F-5/6 No. 32 | A chimeric strain derived from GP388 |
| GP429 | RERE6 Kernel-supply3 gent-gba | The kernel genome fused with the supply3 cassette |
| GP538 | HST08 gent-pir | R6K *pir*-expressing plasmid |
| GP541 | RERE6 kernel-attPv-zeo-oriT::attBv-mLychee-bla | free of helper plasmid |
| GP549 | DH5α λpir | KT-3228 (Biomedal) |
| GP576 | chimera38 | A chimeric strain derived from GP580 |
| GP580 | RERE6 kernel-supply3-(bla::hyg, kan::spec, cat::zeo, kan@D-1/2, cat@J-9/10 pac@D/E gent@I/J) | |
| GP588 | chimera38 pGCORE-star-ΔAvrII | |
| GP595 | hybrid chimera Tet7 strain | The single-genome strain derived from GP588 |

| | | |
|---|---|---|
| GP599 | HST08 pOri2-tet-tra-CV-phiC31int | A conjugal donor strain |
| GP606 | REGE-244A | The pOri2-lacking derivative of GP595 |
| GP609 | RERE6 kernel-supply3-(bla::hyg, kan::spec, cat::zeo, kan@G-9/10) gent-pir pR6K2013-ΔoriT | A conjugal donor strain |
| GP612 | REGE-244 | The plasmid-free derivative of GP606 |
| GP641 | REGE-229 | Two regions of the REGE-244 genome were replaced by the GP609 kernel genome. |

### Table S3. List of plasmids and chromosomes.

| plasmid name | description | sequencing |
|---|---|---|
| pMW118cat-rrnCΔAvrII | *rnnC*(ΔAvrII), *cat* | iSeq 22/07/15 |
| ptRNA100-gltT::zeo | ptRNA100 Δ*gltT* | iSeq 22/07/15 |
| designed kernel genome ver 1.08 | the originally designed kernel genome sequence | only designed |
| W3110&RGF&Kernel | mapping small genomes to wildtype | only designed |
| ForiC2.6 | RCR mini-F vector, *amp* | iSeq (part of others) |
| ForiC2.6_spec_amp | RCR mini-F vector, *amp, spec* | iSeq (part of others) |
| pCP20-Cm | pCP20 Δ*amp* | function-confirmed |
| 2xR1 vector | RCR *ori2* vector, *spec* | iSeq 23/04/21 |
| 2xR2 vector | RCR *ori2* vector, *spec* | iSeq 23/04/21 |
| pOri2bC | RCR *ori2* vector, *kan, amp* | iSeq 23/02/10 |
| pOri2bsC | RCR *ori2* vector, *amp, spec* | iSeq 23/02/25 |
| pMW118 | *amp* | purchased from NIPPON GENE |
| pMW118cat | pMW118 *amp::cat* | iSeq (part of others) |
| pKD46 | ts helper for λ RED recombination, *amp* | ref. Datsenko and Wanner, 2000 |
| pMW118-gba | helper for λ RED recombination, *amp* | ref. Mukai et al., 2020 |
| pMW118-g'ba | helper for λ RED recombination, Δ*gamS*, *amp* | ref. Mukai et al., 2020 |
| gent-gba | helper for λ RED recombination, *gent* | iSeq (part of others) |
| hyg-gba | helper for λ RED recombination, *hyg* | iSeq 24/02/29 |
| hyg-gbaCre | helper for λ RED recombination and Cre-*loxP* recombination, *hyg* | iSeq 24/05/02 |
| hyg-g'baCre | helper for λ RED recombination and Cre-*loxP* recombination, Δ*gamS*, *hyg* | function-confirmed |
| hyg-gba-recA | helper for λ RED recombination, *recA*, *hyg* | iSeq 24/05/02 |
| gent-pir | helper for R6K*ori* replication, *gent* | iSeq 25/06/13 |
| pJExDTT-NoeilR-sfGFP-TtufB_2xAvrII | reporter, *gent* | function-confirmed |
| pTet17-HKint | helper for HK022 integrase, aTc inducible, *gent* | iSeq 24/01/31 |
| pMW118-HKint | helper for HK022 integrase, arabinose inducible, *amp* | iSeq 23/06/16 |
| pMW118-HKxisint | helper for HK022 excisionase and integrase, arabinose inducible, *amp* | function-confirmed |
| dual-int | helper for HK022 and φC31 integrases, arabinose inducible, *amp* | iSeq 23/06/16 |
| pBAD-traRP4min | conjugation *tra* helper plasmid, arabinose-inducible, *amp* | ref. Mukai et al., 2020 |
| pBAD-traRP4min-Plac-Nb3 | conjugation *tra* helper plasmid, arabinose-inducible, *amp* | iSeq 23/04/21 |

| | | |
|---|---|---|
| pRK2013 | conjugation *tra* helper, self-transmissible, *kan* | purchased from Biomedal |
| pTra2013 | conjugation *tra* helper, Δ*oriT*, still self-transmissible, *amp* | iSeq 24/01/31 |
| pR6K2013 | suicide conjugation *tra* helper, self-transmissible, R6K*ori*, *amp* | iSeq 25/01/24 |
| pR6K2013-ΔoriT | suicide conjugation *tra* helper, Δ*oriT*, still self-transmissible, R6K*ori*, *amp* | iSeq 25/07/03 |
| pGCORE-star-ΔAvrII | a chromosomal core, the *ori* region with *oriC**, *kan* | iSeq 21/11/19 |
| pOri2-tet-tra-CV-phiC31int | self-transmissible, helper for φC31 integrase, *tet* | iSeq 25/11/07 |
| gent-recA | helper for RecA, *gent* | iSeq 24/11/20 |
| amp-recA | helper for RecA, *amp* | iSeq 25/09/26 |
| pMWts-tet | ts, *tet* | iSeq 26/01/30 |
| pOri2-tet-recA-2xCV-I-SceI | helper for RecA and I-SceI, *tet* | iSeq 26/03/06 |
| pOri2-2xR2-VHb-sd-ratAB-yadGH | vector for pOri2-ori | iSeq 23/05/19 |
| pOri2-2xR2-ori | for Tus-masking RA-RCR | iSeq 23/06/02 |
| pOri2-2xR1-dif | for Tus-masking RA-RCR | iSeq 23/05/19 |
| pOri2-oridif | core of the kernel genome, clone: 1B-3 | iSeq 23/06/16 |
| pOri2-2xR1-MC2fst | for Tus-masking RA-RCR | iSeq 23/04/28 |
| pOri2-2xR2-MC2snd | for Tus-masking RA-RCR | iSeq 23/04/28 |
| pOri2-2xR1-MC3fst | for Tus-masking RA-RCR | iSeq 23/04/28 |
| pOri2-2xR2-MC3snd | for Tus-masking RA-RCR | iSeq 23/04/28 |
| pOri2-2xR1-MC4 | for Tus-masking RA-RCR | iSeq 23/05/19 |
| pOri2-2xR2-MC5 | for Tus-masking RA-RCR | iSeq 23/05/19 |
| dMC1 on ForiC2.6specamp | MC1, ΔoriC2.6, clone: 8E8 1-1-1 | iSeq 23/05/19 |
| dMC2 on 2xterAvrII | MC2, ΔoriC2.6, clone: 1D1 | iSeq 23/06/02 |
| dMC3 on 2xterAvrII | MC3, ΔoriC2.6, clone: 4D2-1 | iSeq 23/06/16 |
| dMC4 on ForiC2.6 | MC4, ΔoriC2.6, clone: 1F2-2 | iSeq 23/03/31 |
| dMC5 on ForiC2.6specamp | MC5, ΔoriC2.6, clone: 2H2-1 | iSeq 23/04/21 |
| pOri2kan-attBcc-attP-spec-oriT-attPcc | mobile vector for MC2 | not sequenced |
| pOri2kan-attBcc-attPv-zeo-oriT-attPcc | mobile vector for MC4 | iSeq (part of others) |
| MC2_for_conjugation | transferrable, excisable, integrative | not sequenced |
| MC4_for_conjugation | transferrable, excisable, integrative, clone: i-2C7 | iSeq 23/07/21 |
| MC2-oridif | MC2 was fused into pOri2-oridif, clone: a | iSeq 23/08/04 |
| MC4-oridif | MC4 was fused into pOri2-oridif | iSeq 23/08/04 |
| MC3_for_oriT-POP | transferrable payload, integrative | not sequenced |

| | | |
|---|---|---|
| MC1a_for_oriT-POP | transferrable payload, integrative, clone #3 | iSeq 24/02/29 |
| MC1b_for_oriT-POP | transferrable payload, integrative | iSeq 24/02/29 |
| MC4_for_oriT-POP | transferrable payload, integrative, clone #2 | not sequenced |
| MC5_2xattPv_for_oriT-POP | transferrable payload, integrative using two *attPv* sites | not sequenced |
| MC2-oridif-MC3 | MC3 was fused into MC2-oridif, clone: small2 | iSeq manual 23/09/28, 29 |
| MC2-oridif-MC3-attB-attBv | for the next round of oriT-POP cloning | not sequenced |
| MC2-oridif-MC3-MC4 | MC4 was fused into MC2-oridif-MC3 | not sequenced |
| MC2-oridif-MC3-MC4-attB-attBv | for the next round of oriT-POP cloning | not sequenced |
| MC1b_MC2-oridif-MC3_MC4 | MC1b was fused into MC2-oridif-MC3-MC4, clone: 14-24-8 | iSeq manual 24/03/13 |
| 1.7Mb_MC1to5 | The original kernel genome | iSeq manual 24/10/02 |
| 1.7Mb_mLychee | kernel with mLychee | iSeq manual 25/08/08 |
| pFts-Supply3 | mini-F, ts, supply3-*kan* for RMCE | iSeq 25/02/14 |
| 1.7Mb_supply3 | kernel with mLychee and supply3 | not sequenced |
| 1.7Mb_hygzeospecgentpaccatstarkanstar | kernel with mLychee, supply3, and multiple markers (* 2x I-SceI sites) | not sequenced |

### Table S4. List of DNA cassettes.

| cassette name | description |
|---|---|
| cat(wildtype) | |
| cat(enhanced) | promoter-enhanced |
| cat* | inserted between J-9/10 on the kernel genome, one I-SceI site |
| kan | promoter-enhanced |
| kan* | inserted between D-1/2 on the kernel genome, one I-SceI site |
| spec | promoter-enhanced |
| zeo | *shble* |
| tet | for *endA1* knockout (ref. Fujita *et al*., 2022) |
| hyg | to replace the *bla* on the kernel genome |
| pac | inserted between D/E on the kernel genome |
| Plac-gent | inserted between I/J on the kernel genome |
| attR-cat | λ RED setup to regenerate *attB* |
| attRv-cat | λ RED setup to regenerate *attBv* |
| kan-attB | λ RED setup to insert *attB* |
| spec-oriT | for oriT-POP (start) |
| zeo-oriT | for oriT-POP (start) |
| attPv-zeo-oriT | for oriT-POP (stop) |
| attP-spec-oriT | for oriT-POP (stop) |
| attPv-bsdbsd | the 2nd *attPv* inserted between the *sspB/ytfL* of MC5 for oriT-POP |
| attB-loxP-kan | inserted @*torT* of the 1.3-Mb-fused genome to replace the *zeo-oriT* |
| pac-loxP | inserted @*torS* of the 1.3-Mb-fused genome to replace the *attL-spec-oriT* |
| attB-oriC-attP | the *attB-oriC-attP* locus |
| SUE11727 | oligo DNA for *gyrA96* reversion to wildtype |
| CV-avrII@rhaBS | for developing RERE6 (marker-removed) |
| CV-avrII@melRA | for developing RERE6 (marker-removed) |
| CV-avrII@uxaC-exuT | for developing RERE6 (marker-removed) |
| aTc-avrII@bscRE | for developing RERE6 (marker-removed) |
| aTc-avrII@ttdRA | for developing RERE6 (marker-removed) |
| aTc-avrII@ulaGA | for developing RERE6 (marker-removed) |

| | |
|---|---|
| lox66-kan-lox71 | to replace the *spec* gene next to the *oriT* on the kernel genome |
| lox71-zeo-loxP25766 | to replace the *loxP* site on the kernel genome |
| mLychee-bla | to replace the *oriT-zeo-attPv* on the kernel genome |

### Table S5. List of NGS data.

| strain name | strain number | date folder | file name |
|---|---|---|---|
| SQ171/pMW118cat-rrnCΔAvrII/ptRNA100-gltT::zeo | GP207 | 20220715 | 2_S2 |
| 1.3 Mb pTra2013 pTet17-HKint | XGP_13p | 20240313 | 2_S2 |
| 1.3 Mb with cat in genome pTra2013 hyg-gba | XGP_13g | 20240313 | 3_S3 |
| 1.7 Mb in genome | XGP_17 | 20240821 | 30_S30 |
| HST08 ΔattB endA1::tet Kernel pMW118-g'ba | GP346 | 20241002 | 2_S2 |
| chimera_8 | XGP_8 | 20241002 | 4_S4 |
| chimera_24 | XGP_24 | 20241002 | 6_S6 |
| chimera_407-1 | GP365 | 20241018 | 4_S4 |
| chimera_407-2 | GP366 | 20241018 | 5_S5 |
| chimera_G/H-1 | GP372 | 20241018 | 8_S8 |
| chimera_G/H-2 | GP373 | 20241018 | 9_S9 |
| chimera_388-e | GP375 | 20241029 | 32_S32 |
| chimera_I/Cys-old | GP377 | 20241029 | 34_S34 |
| chimera_I/Cys-new-a | GP378 | 20241029 | 35_S35 |
| chimera_I/Cys-u | GP379 | 20241029 | 36_S36 |
| chimera_H-7/8 No. 4 | GP380 | 20241029 | 37_S37 |
| chimera_H-7/8 No. 7 | GP381 | 20241029 | 38_S38 |
| chimera_C-5/6 No. 13 | GP389 | 20241120 | 24_S24 |
| chimera_F-5/6 No. 22 | GP390 | 20241120 | 25_S25 |
| chimera_F-5/6 No. 32 | GP391 | 20241120 | 26_S26 |
| RERE6 Kernel-mLychee | GP541 | 20250808 | 2_S2 |
| chimera38 | GP576 | 20250926 | 70_S70 |
| REGE-244A | GP606 | 20251219 | 1_S1 |
| REGE-229 | GP641 | 20260424 | 3_S3 |

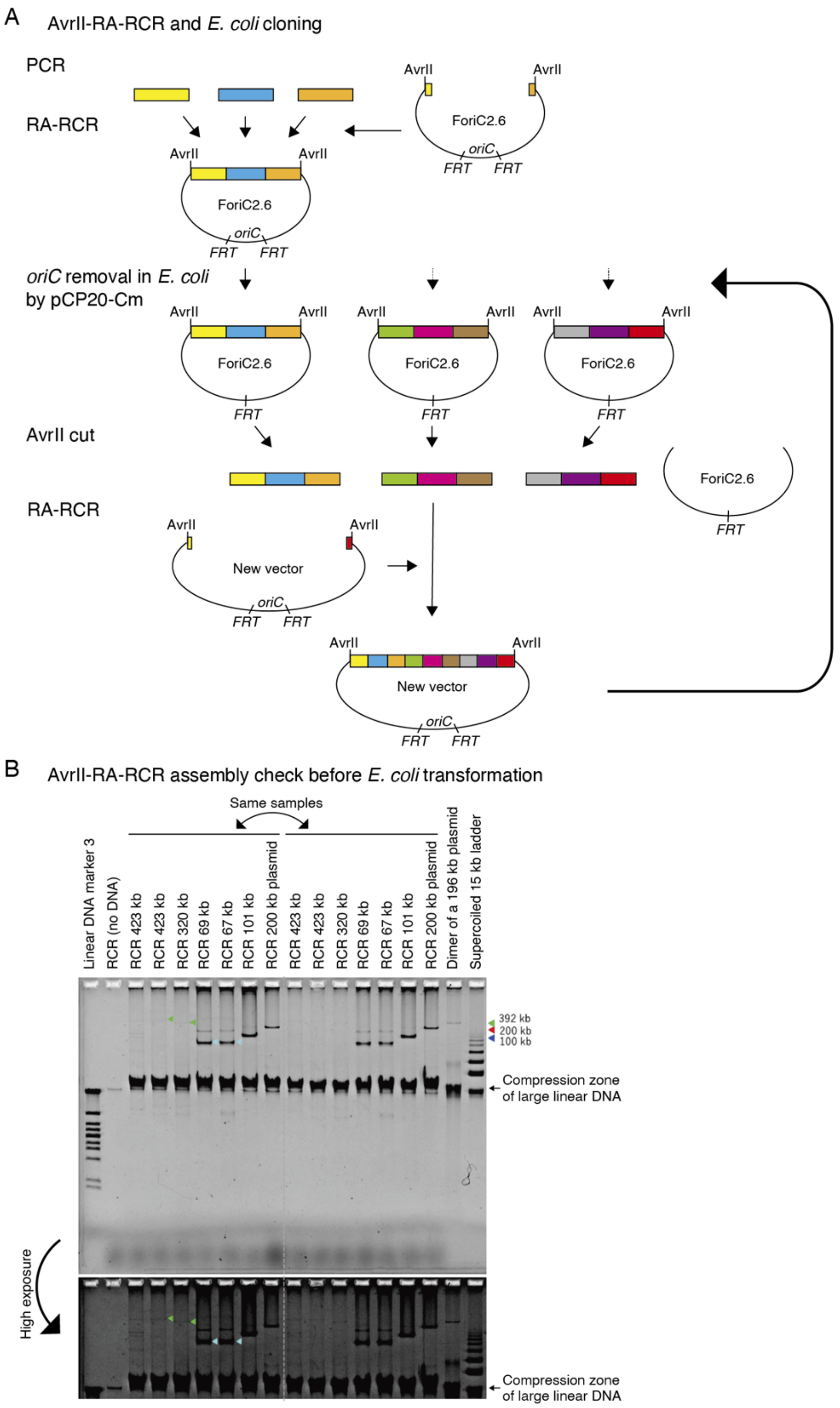


**Figure S1. AvrII-RA-RCR method.** The procedure **(A)** and an example **(B)** are shown. Supercoiled plasmids were analyzed by canonical agarose gel electrophoresis. Some bands are indicated with arrows.

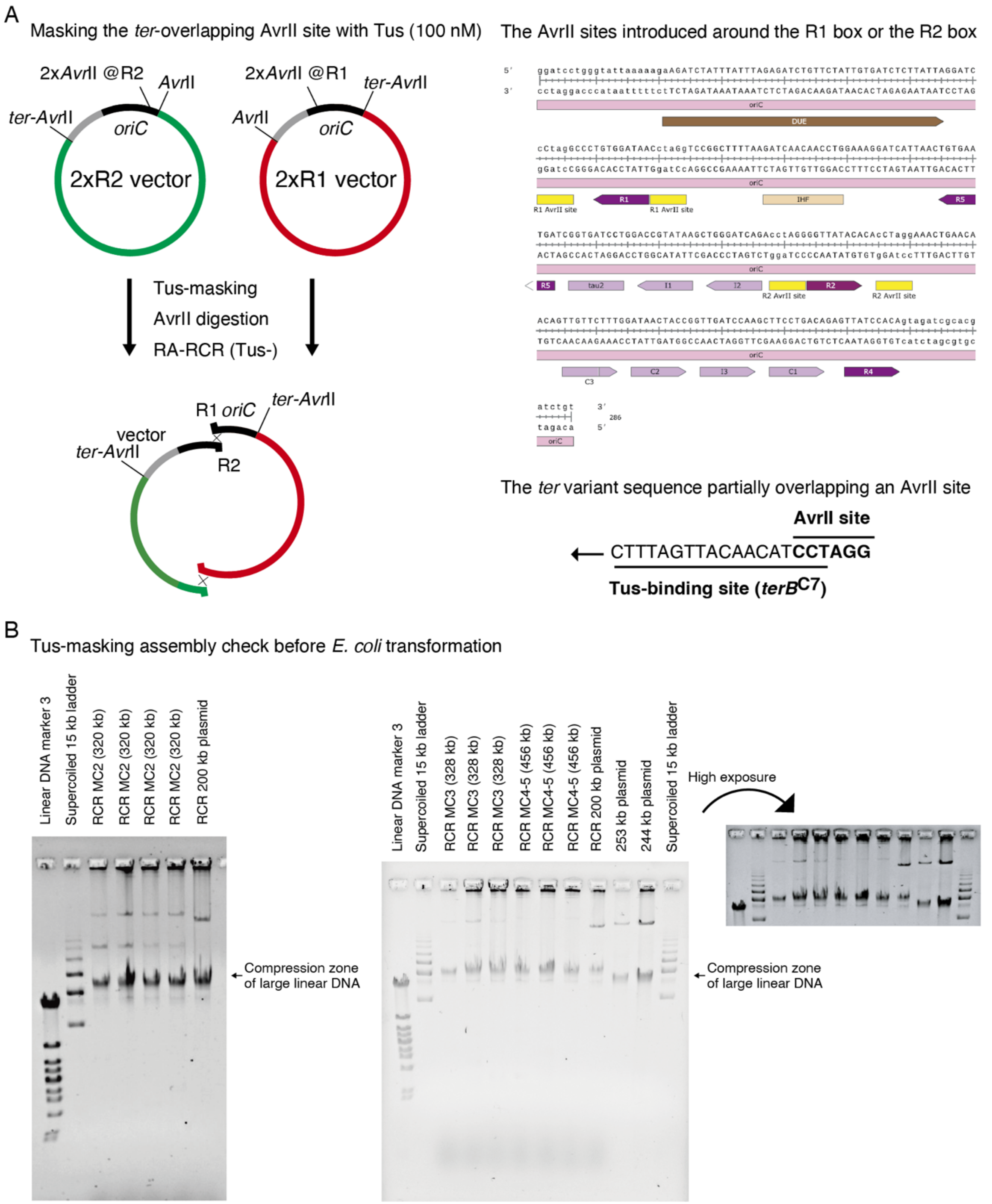


**Figure S2. Tus-masking assembly method.** The procedure **(A)** and examples **(B)** are shown. Supercoiled plasmids were analyzed by canonical agarose gel electrophoresis. The MC4-5 megachunk was not successfully cloned in *E. coli* cells.

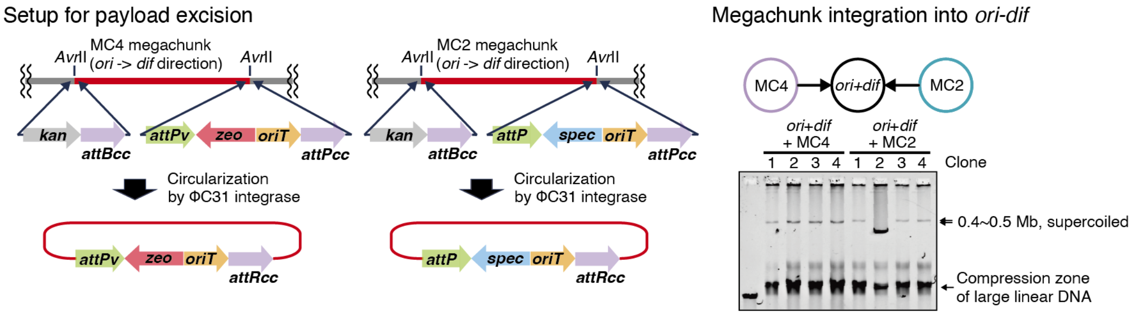


**Figure S3. Megachunk payload fusion with the *ori-dif* core using two integrase systems.** The MC4 and MC2 megachunk regions were flanked by the φC31 *attB*$^{cc}$ and *attP*$^{cc}$ variant sequences on the *ori2* plasmid vector. After their conjugal transfer, the megachunk payload was excised via circularization from the vector by φC31 integrase and integrated into the *ori-dif* core by HK022 integrase. *E. coli* cells harboring the fusion chromosomes were isolated by antibiotic selection. The fusion chromosomes were purified by miniprep and resolved by agarose gel electrophoresis analysis.

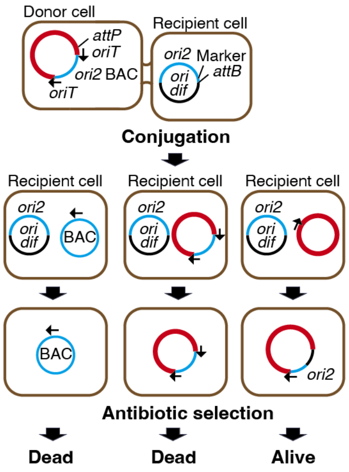


**Figure S4. Plasmid payload delivery.** Scheme of the megachunk fusion to the core plasmid by plasmid payload delivery via conjugation. The *oriT*-flanked megachunk payload on an *ori2* BAC vector can be conjugally transferred from the donor cells, circularized at *oriT*, and site-specifically integrated into the core chromosome in the recipient cells. Since two *ori2* plasmids cannot be maintained in the same cell (plasmid incompatibility), antibiotic selection of the recipient *ori2* plasmid eliminates cells harboring the donor *ori2* plasmid. For simplicity, several patterns are omitted from the picture.

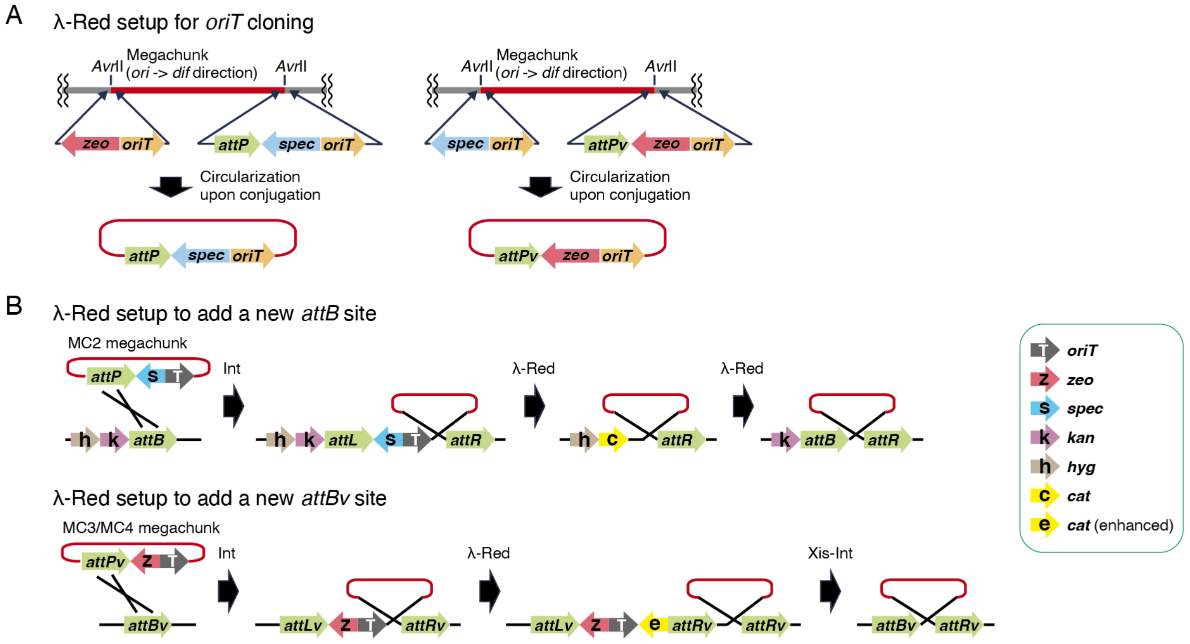


**Figure S5. Setup for *oriT*-POP cloning via λ Red recombination. (A)** Each megachunk cloned on an *ori2* vector was flanked by the indicated DNA cassettes while removing the AvrII sites. **(B)** When needed, a new *attB* or *attBv* sequence was introduced via λ Red recombination. The HK022 integrase (Int) mediates integration, while it mediates excision in the presence of the excisionase (Xis). To facilitate antibiotic selection, the *cat* promoter was enhanced.

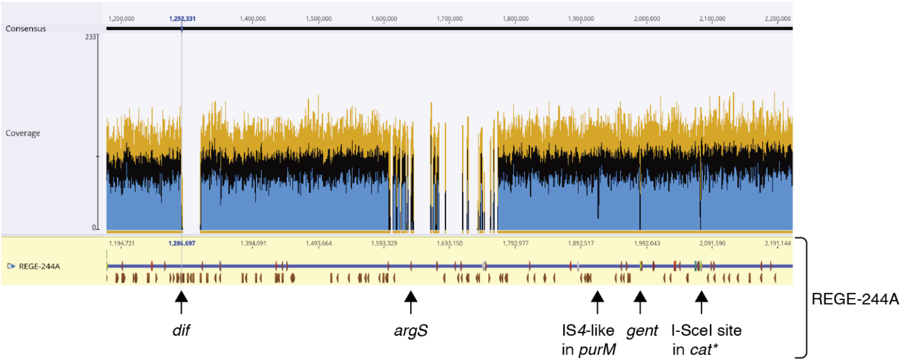


**Figure S6. Mapping the NGS reads of REGE-229 to the REGE-244A map.**

### References

1. Datsenko, K.A. and Wanner, B.L. (2000) One-step inactivation of chromosomal genes in *Escherichia coli* K-12 using PCR products. *Proc Natl Acad Sci U S A* **97**, 6640-6645.

2. Hirokawa, Y., Kawano, H., Tanaka-Masuda, K., Nakamura, N., Nakagawa, A., Ito, M., Mori, H., Oshima, T. and Ogasawara, N. (2013) Genetic manipulations restored the growth fitness of reduced-genome *Escherichia coli*. *J Biosci Bioeng*, **116**, 52-58.

3. Mukai, T., Yoneji, T., Yamada, K., Fujita, H., Nara, S. and Su'etsugu, M. (2020) Overcoming the Challenges of Megabase-Sized Plasmid Construction in *Escherichia coli*. *ACS Synthetic Biology*, **9**, 1315-1327.

4. Yoneji, T., Fujita, H., Mukai, T. and Su'etsugu, M. (2021) Grand scale genome manipulation via chromosome swapping in *Escherichia coli* programmed by three one megabase chromosomes. *Nucleic Acids Research*, **49**, 8407-8418.

5. Fujita, H., Osaku, A., Sakane, Y., Yoshida, K., Yamada, K., Nara, S., Mukai, T. and Su'etsugu, M. (2022) Enzymatic Supercoiling of Bacterial Chromosomes Facilitates Genome Manipulation. *ACS Synthetic Biology*, **11**, 3088-3099.
